# Not All Charges Are Equal: Side-Chain Chemistry Reshapes the Disordered Ensemble of *α*-synuclein

**DOI:** 10.64898/2026.06.07.730657

**Authors:** Gil Koren, Nuno P. Fernandes, Asaf Sivan, Rawan Aodeh, Dean Yona, Mathar Kravikass, Yuval Keidar, Kresten Lindorff-Larsen, Roy Beck

**Author notes:** Author contributions: G.K. and R.B. designed research; G.K. purified proteins with the assistance of R.A. and Y.K.; G.K., D.Y.and M.K. preformed SAXS measurements and analysis; A.S.and N.P.F. performed and analyzed molecular dynamics simulations with guidance from K.L.-L.; G.K and R.B. wrote the paper. K.L.-L. holds stock options in and is a consultant for Peptone Ltd. The remaining authors declare no competing interest.

## Abstract

The conformational ensembles of intrinsically disordered proteins (IDPs) are governed by the balance of electrostatic and hydrophobic interactions encoded in their primary sequences. Current polymer-physics models of IDPs frequently group amino acids by coarse-grained properties, such as net charge, often overlooking the distinct residues’ side-chain chemistries. Analysis across the IDP database reveals that these sequences are sensitive to specific residue identities, where the substitution of aromatic, proline, or hydrophobic groups serves as a primary driver of chain dimensions. However, these data also highlight that even subtle chemical variations between similarly charged residues can consistently shift global compaction. Here, we explore the role of residue identity using small angle X-ray scattering (SAXS) of seven *α*-synuclein variants with progressively increasing numbers of lysine-to-arginine substitutions, two positively charged amino acids with different side-chain chemistry. We show that increasing arginine content drives a systematic compaction of the conformational ensemble, although variants with identical number of substitutions but different positional arrangements suggest influence to the sequence context. Moreover, while increasing salt concentrations shift the structural ensemble from a Gaussian toward a self-avoiding-walk statistics, the arginine-dependent contraction trend remains robust across both regimes. Molecular-dynamics simulations combined with SAXS data reveal that arginine substitutions reduce ensemble heterogeneity by stabilizing transient long-range contacts. Finally, aggregation assays demonstrate that this arginine-driven compaction correlates with an accelerated transition to amyloid fibrils. Our findings demonstrate that chemically subtle substitutions between similarly charged residues can fundamentally reshape the conformational ensemble of IDPs, suggesting that side-chain identity is a critical, yet underappreciated, determinant of protein disorder and proteotoxicity.

Significance Statement
Intrinsically disordered proteins (IDPs) are critical to cellular signaling and neurodegeneration, yet our ability to predict their behavior remains limited by a “coarse-grained” understanding of their sequences. We expand the view that net charge and patterning is a primary determinant of IDP dimensions by showing that lysine and arginine, residues identical in charge, exert opposite effects on the conformational landscape of *α*-synuclein model-system. We find that arginine substitutions act as a “molecular glue”, driving protein compaction and reducing ensemble heterogeneity. Crucially, this compaction accelerates amyloid aggregation, overturning the conventional assumption that collapsed states protect against fibrillization. This work demonstrates that side-chain identity and patterning are vital for protein homeostasis, providing a new framework for the rational design of IDP-based therapeutics.

---

Intrinsically disordered proteins (IDPs) lack a stable three-dimensional structure under physiological conditions, yet they play crucial roles in numerous cellular processes, including signaling, regulation, and phase separation (1). Similar to folded proteins, IDPs are sensitive to sequence composition and patterning, which govern their conformational ensembles and functional interactions (2–4). However, because IDPs are highly dynamic, heterogeneous, and context-dependent, understanding and predicting their conformational ensembles remains a fundamental challenge in structural biology (5, 6).

The conformational ensembles of IDPs are typically characterized by mean-field apparent scaling factors (*ν*) and probability distributions of global parameters such as the radius of gyration (*R*_*g*_). Indeed, previous studies have shown that these global properties vary systematically with sequence-encoded electrostatics, including the net charge per residue (NCPR) and the fraction of charged residues (FCR) (7–10). Moreover, correlations between IDP charge content and physical dimensions can be observed not only across different proteins but also within distinct segments of the same protein (11).

As a first-order descriptor, NCPR has proven useful in predicting ensemble expansion or compaction across many IDPs. However, NCPR is a global average and thus is inherently limited: it cannot capture sequence heterogeneity or the effects of charge distribution, which are critical for understanding the behavior of sequences that contain both positively and negatively charged residues (polyampholytic) or are rich in polar amino acids. In such cases, a deeper understanding requires the consideration of higher-order descriptors, such as charge patterning (3, 12, 13) and the presence of hydrophobic residues (14–16).

Analyses of experiments and simulations have shown that variations in the chain dimensions across diverse IDPs are better explained when accounting for additional sequence features such as charge clustering, polarity patterning, and glycine content, rather than NCPR alone (10, 17). Further complexity arises from charge regulation, where charges are not fixed but depend on the local environment and interactions with counterions from the electrolyte (18). Dipole formation is conformation dependent, while the conformation itself is influenced by the distribution of charges and dipoles along the chain (19). Importantly, even residues with the same nominal charge can differ significantly in their side-chain structure, hydrogen bonding potential, and hydrophobicity (20), leading to sequence-specific effects that are not captured by coarse metrics like NCPR. Thus, to fully understand the sequence–ensemble relationship in IDPs, one should consider not just the quantity of charge, but also its type and the relevant chemical context.

Analysis of UniProt entries across disordered domains reveals that arginine is the most frequent amino acid involved in disease-associated point mutations (Fig. S1). This enrichment suggests that specific charged residues, particularly arginine, play a disproportionate role in maintaining the structural and functional balance of intrinsically disordered regions. This finding is surprising, as such fine-tuned sequence modulation is typically associated with structured proteins, whereas IDPs are often thought to be governed by more coarse-grained sequence–function relationships as described above.

Roesgaard et al. (21) used a combination of NMR, small angle X-ray Scattering (SAXS), simulations, biochemistry and cell biology to study differential effects of aspartic (D) and glutamic (E) acid in the protein DSS1. Sequence analyses show E is more enriched in IDPs than D, yet the origins of this remains unknown. Experiments probing global and functional properties in DSS1 revealed only small differences in variants with all-D, all-E or D and E swapped (21). In contrast, NMR experiments revealed substantial differences between D and E, likely due to differences in stabilizing helical structures (21).

As another example, lysine (K) and arginine (R) both carry a charge of + 1 at physiological pH, yet they are fundamentally different in their side-chain chemistry, hydration properties, and potential for specific interactions (20, 22). Arginine binds negatively charged amino acids with higher affinity than lysine (23), indicating that positive charges in proteins should not be considered equivalent when carried by lysine versus arginine. Moreover, while both cations can engage in cation–*π* interactions, these interactions are stronger for arginine due to its guanidinium group (24). Arginine alone can further engage in *π* − *π* interactions via its guanidium group as well as in multiple hydrogen bonding interactions (25, 26). Its ability to remain protonated even in unfavorable environments arises from this same group establishing extensive hydrogen bonding with polar protein atoms and bound water (27). Thus, the free energy of hydration obtained for arginine is consistently smaller than that for lysine (20), and guanidinium ions self-associate more strongly than ammonium (28). Furthermore, the positive heat capacity of hydration is ∼ 2.3 times greater for arginine compared to lysine (20).

Consistent with these physicochemical differences, arginine-rich sequences are expected to adopt more globular conformations, whereas lysine-rich sequences should behave more like self-avoiding walks (29). This mechanism is further supported by experimental measurements that arginine rich polyampholytic IDPs tend to adopt significantly more compact conformations than their lysine rich counterparts (30) and that lysine-to-arginine substitutions in the low complexity domain of hnRNPA1 makes the protein more compact (31). Arginine-rich peptides self-associate more strongly than corresponding lysine-containing peptides (32), and arginine residues tend to increase the lifetimes of interchain contacts at low salt concentrations (33). These distinctions contribute to the differential roles of arginines versus lysines in driving phase separation (22, 31, 34, 35).

Computational approaches, including molecular dynamics (MD) simulations, including with coarse-grained models, and polymer physics-based frameworks, have significantly advanced our ability to describe IDP conformational ensemble behavior (36). However, these models often rely on simplifying assumptions about residue–residue interactions and solvation effects, and thus struggle to resolve subtle, sequence-specific experimental differences. For example, an early data-driven coarse-grained model for IDPs represented arginine as more interaction prone than lysine (37), whereas a later model represented arginine as the least interaction prone of all 20 amino acids (38).

More recent coarse-grained models, such as CALVADOS (39), HPS-Urry (40) and Mpipi (41) show increased interactions with arginine relative to lysine, though differ in the relative strengths of interactions. Simulations with these types of models can also be used to train machine learning models for studying IDPs, taking into account the amino-acid specific flavor (17, 42–46). For example, ALBATROSS (44) can estimate average global properties of disordered proteins from sequence alone, including *R*_*g*_, *ν*, and asphericity. By training neural networks on large ensembles of coarse-grained simulations to directly learn the mapping between sequence features and ensemble-averaged structural descriptors, STARLING (45), as another example, further predicts average inter-residue distance maps, thus enabling rapid and high-resolution comparison of subtle conformational modulation resulting from sequence modifications.

The *α*-synuclein (*α*Syn) protein is an ideal model system for studying how charge identity and patterning influence the conformational properties of IDPs. Its primary sequence displays pronounced heterogeneity, including clusters of positively charged residues, hydrophobic motifs, and highly acidic regions, all embedded within a fully disordered backbone (47, 48). Notably, *α*Syn contains 15 lysine residues but no arginine in its native sequence, making it particularly well suited for dissecting the role of sequence-encoded interactions, such as long-range electrostatics and side-chain specific effects.

Beyond its value as a biophysical model, *α*Syn is also prone to aggregation, and its misfolding into amyloid fibrils is a central pathological feature of Parkinson’s disease (49). Mechanistically, aggregation is thought to proceed via a nucleation-dependent process, in which long-range intramolecular interactions between the N- and C-terminal domains shield the hydrophobic non-amyloid-*β* component (NAC) region, thereby regulating aggregation kinetics (50). Disruption of this shielding can expose the NAC region and promote intermolecular interactions that drive fibril formation. However, some studies suggest the opposite, that the N- and C-terminal regions do not effectively shield the NAC domain (51). In addition, several familial Parkinson’s disease–associated mutations in *α*Syn, such as A30P, E46K, and A53T, have been shown to accelerate the aggregation process and enhance fibril formation (52, 53). A recent study systematically probed lysine-to-glutamine substitutions on *α*Syn aggregation and showed that charge explains some but not all of the observed effects (54). Despite extensive investigation, the precise relationships between conformational ensemble features, sequence-level perturbations, and aggregation propensity remain unresolved.

In this study, we experimentally investigate how substitutions of lysine to arginine alters the conformational ensemble of *α*Syn. To experimentally assess conformational changes induced by lysine-to-arginine substitutions, we employed SAXS to measure global dimensions and flexibility in solution (11, 55). Furthermore, we combined molecular dynamics (MD) simulations with the Ensemble Optimization Method (EOM) (56, 57) and Bayesian/Maximum Entropy (BME) reweighting (58) to extract the underlying structural distributions consistent with the SAXS data, enabling a detailed description of both average conformational trends and ensemble heterogeneity. Our results reveal an arginine-induced compaction of the ensemble, which persists across varying salt conditions. Moreover, we uncover a dependence on arginine sequence heterogeneity, underscoring the importance of the local sequence context beyond overall composition (21). These findings show that even residues sharing the same charge can give rise to distinct structural outcomes.

## Results

### *α*-Synuclein Model System

We choose *α*Syn as an IDP model due to its fully disordered nature in solution and the absence of arginine residue in its native sequence. The *α*Syn sequence consists of three distinct domains (Fig. 1A) (48). The N-terminal lipid binding domain (residues 1–60) is dominated by five consensus sequences (59) K(A\T)K(Q\E)(Q\G)V which make this domain a basic polyelectrolyte with FCR = 0.25 and NCPR = 0.067. This domain interacts with lipid membranes and adopt two *α*-helices upon binding (48, 60). The non-amyloid-*β* component (NAC) domain (residues 61–95) is largely hydrophobic with minimal charges (FCR = 0.086), and is involved in protein aggregation. Lastly, the C-terminal domain (residues 96–140) is highly acidic (FCR = 0.4, NCPR = -0.267) and enriched in proline residues.

**Fig. 1.**
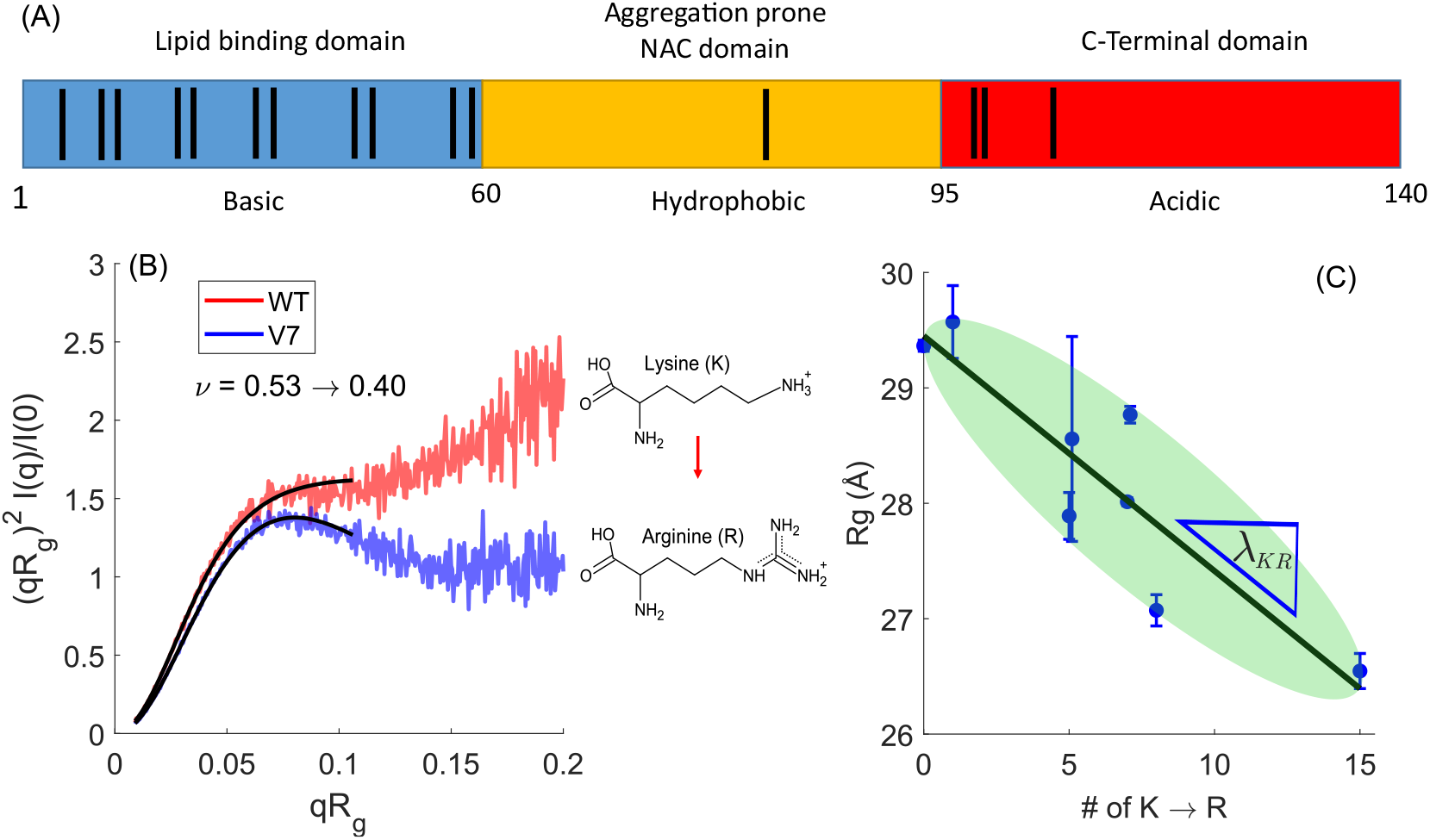
Lysine-to-arginine (K→R) substitutions tune *α*Syn compaction. (A) Schematic of the *α*Syn sequence, showing the N-terminal lipid-binding domain (residues 1–60, blue), the NAC domain (residues 61–95, yellow), and the acidic C-terminal domain (residues 96–140, red). Lysines are indicated by vertical black bars. (B) Dimensionless Kratky plot of WT (red) and V7 (blue) variant, where all 15 lysine (K) are substituted to arginine (R) residues. Solid black lines represent the Molecular Form Factor (MFF) fits. Inset: Change of the apparent scaling exponent (*ν*) for the transition from WT to V7. (C) Radius of gyration (*R*_*g*_) measured by SAXS decreases with increasing lysine-to-arginine (K→R) substitutions. The slope of this correlation, *λ*_KR_, quantifies the compaction effect of K→R substitutions. The green area indicates (*R*_*g*_) values for all possible K→R positional variants of *α*Syn predicted by ALBATROSS, illustrating the range of conformations.Errors are calculated as explained in the Methods section.

To investigate how the identity of positively charged residues influences the conformational ensemble of *α*Syn, we generated a panel of seven variants carrying increasing numbers of lysine-to-arginine (K→R) substitutions (Table 1, Supplementary Table S1). These K→R substitutions are expected to alter the hydrogen bonding network, as arginine’s guanidinium group allows for greater charge delocalization and the formation of more extensive hydrogen bonds (Fig. 1 inset).

**Table 1.**
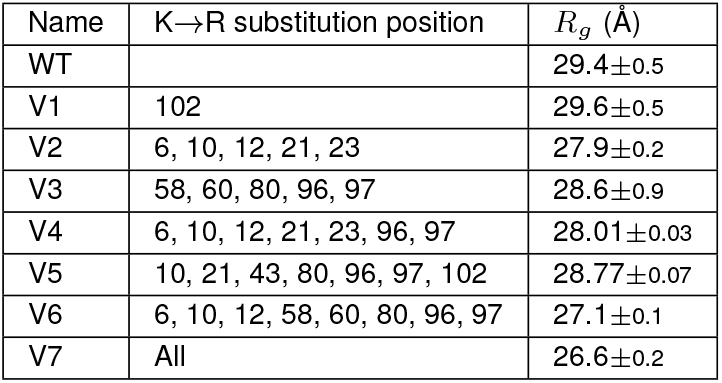
Variants of *α*-Synuclein analyzed in this study, indicating the positions of K→R substitutions and the corresponding *R*_*g*_ measured under salt-free conditions. Errors are calculated as explained in the Methods section.

We expressed and purified all variants from *E*.*coli* and confirmed their disordered nature using circular dichroism (CD, see Fig. S2) and SAXS. CD secondary structure analysis (61) revealed an increase of 15% *β*-sheet content for the V7 mutant, in which all 15 lysines are replaced with arginines, whereas the WT and variants V1–V6 exhibit less than 5% *β*-sheet content. SAXS Kratky analysis revealed profiles characteristic of IDPs for all constructs. Notably, all variants display highly similar dimensionless Kratky curves to WT with an increase profile at high scattering angles (Fig. S3), indicating that the overall disordered nature and global conformational behavior are preserved across the series regardless of the K→R substitutions.

### Substitutions Across All IDPs are Side-chains Specific

To evaluate the structural impact of K→R substitutions we employed SAXS to measure the *R*_*g*_ of each variant in solution. Scattering intensity curves were collected at three different protein concentrations to assess possible protein aggregation.

To quantify the chain dimensions, we performed Molecular Form Factor (MFF) analysis (62) to extract the apparent polymer scaling exponent *ν*. This analysis revealed a decrease in *ν* from 0.53 for WT to 0.4 for the V7 mutant, indicating a transition toward increased chain compaction upon full *K* → *R* substitution (Fig. 1B).

Consistent with this trend, *R*_*g*_ values progressively decreased with increasing numbers of K→R substitutions (Fig. 1C and Table 1), suggesting a systematic compaction of the conformational ensemble. While individual variants showed some variability, a clear linear negative trend was observed, with the fully substituted variant (*i*.*e*., K→R at all 15 positions) exhibiting the most compact ensemble. The inverse relationship indicates that arginine substitutions, despite retaining the same net positive charge, reduce the overall dimensions of the ensemble compared to lysine.

We also found that two variants with the same number of substitutions (V4 and V5, both with 7 K→R substitutions) may have subtly different *R*_*g*_ (Fig. 1C); a similar difference is observed between variants V2 and V3, which both contain 5 K→R substitutions, although here the variation is within the experimental error range. These differences indicate that the global trend of arginine-induced compaction may be modulated by sequence heterogeneity and context, where the specific positioning of arginine residues within the chain influences the resulting conformational ensemble.

To further investigate the K→R substitutions results, we employed the ALBATROSS computational predictor, a deep learning model trained to estimate *R*_*g*_ for IDPs (44). The ALBATROSS predictions (Fig. 1C, green ellipse) represent the range of possible *R*_*g*_ values for all potential K→R substitution combinations. While the total range predicted by the model aligns well with the spread of our experimental SAXS data, the model does not capture all of the fine-grained differences between specific positional isomers. For instance, ALBATROSS does replicate the experimental observation that at 150 mM NaCl *R*_*g*_ (*V* 2) *> R*_*g*_ (*V* 3) but not the observation that *R*_*g*_ (*V* 5) *> R*_*g*_ (*V* 4) (Fig. S4). This suggests that while the model effectively identifies the overall impact of side-chain chemistry on chain dimensions, it may lack the sensitivity to resolve how specific sequence patterns and local contexts fine-tune the conformational landscape of *α*Syn. We do note that these differences are small within the experimental error range when analyzing with alternative methods (Supplementary Table S2).

To quantify the relationship between lysine-to-arginine substitutions and global chain dimensions, we define *λ*_*KR*_ as the slope between changes in radius of gyration *R*_*g*_ and the number of K→R substitutions. Following, *λ*_*KR*_ is normalized to the WT radius of gyration 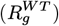 to account for proteins of different lengths (Fig. 2A). Using ALBATROSS, and CALVADOS molecular dynamics simulations we analyzed all IDP sequences from the DisProt database (63). This predictive analysis reveals a similar trend for K→R substitutions, all showing compaction with negative values for *λ*_*KR*_. In fact, the value of |*λ*_*KR*_| for *α*Syn is among the smallest in the distribution of *λ*_*KR*_ values across all IDPs. Moreover, *R*_*g*_ scaling analysis across the entire dataset with substituting all K to R and all R to K, show a systematic change from random coil (all R, *ν* = 0.51) to self-avoiding walk (all K, *ν* = 0.57) statistics (Fig. S5).

**Fig. 2.**
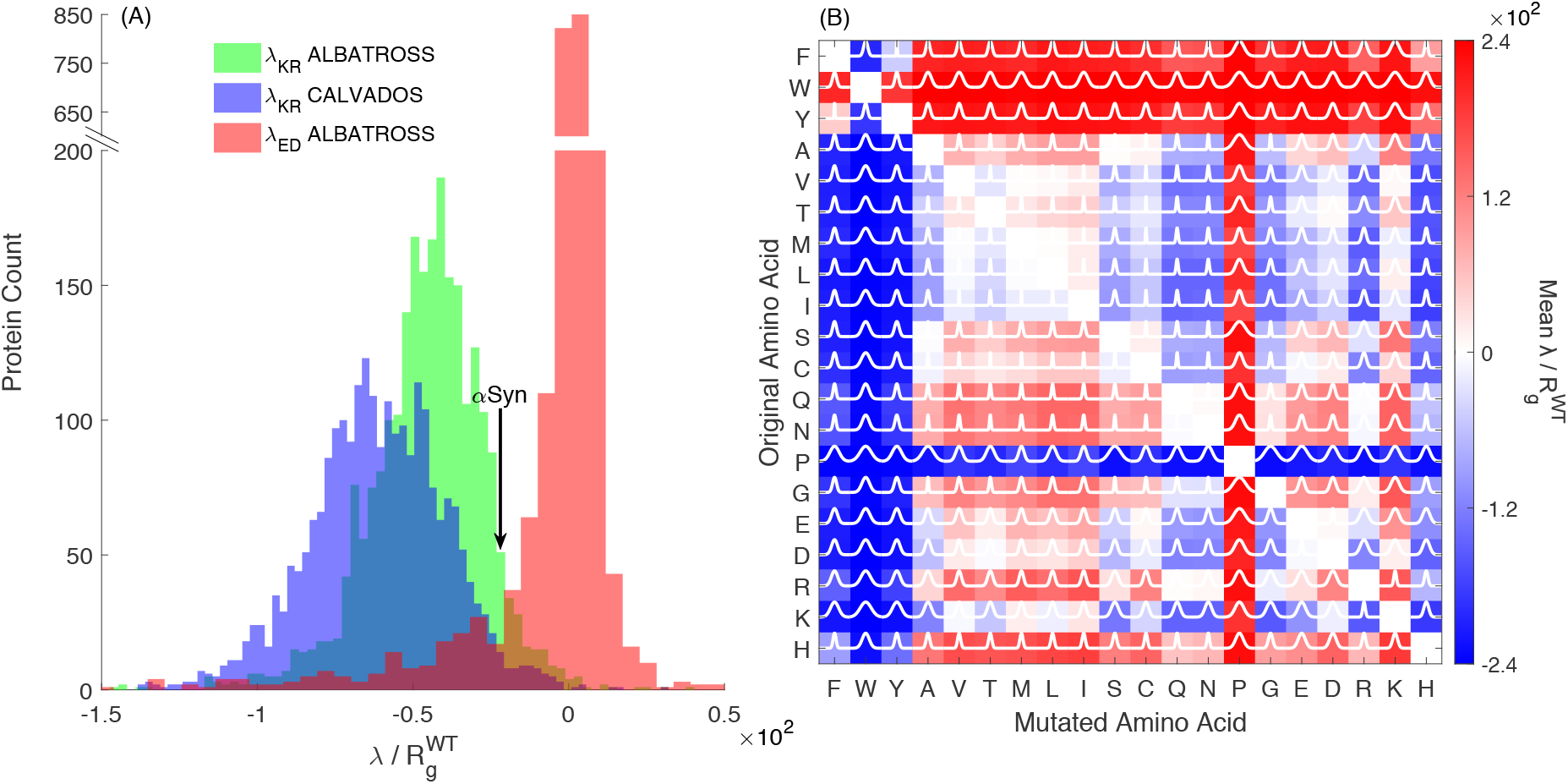
(A) Histograms of ALBATROSS *λ*_KR_ (green), CALVADOS *λ*_KR_ (blue) and ALBATROSS *λ*_ED_ (red) values, normalized by the WT protein’s *R*_*g*_, across all DisProt sequences. The 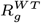 normalization accounts for variations in proteins’ length across the dababase. Black arrow indicate the *α*Syn values for K→R mutations from SAXS. The results highlight the protein’s relative sensitivity to charge-conserving substitutions among intrinsically disordered proteins. (B) 2D heatmap showing the predicted effects of amino acid substitutions on sequence properties, calculated using ALBATROSS. Each cell represents a substitution from the original amino acid (rows) to the target amino acid (columns). The color indicates the scaled predicted mean *λ* value, with blue for negative and yellow for positive values. Overlayed white curves show Gaussian fits for each cell.

We compared these basic substitutions (K→R) to their acidic counterparts (E→D) (Fig. 2A). Unlike the K→R transitions, the distribution for E→D substitutions (*i*.*e., λ*_*ED*_) is narrow and centered around zero. This suggests that acidic residues are largely interchangeable regarding global chain dimensions, whereas the substitution of lysine with arginine introduces a distinct, systematic compaction. These observations are in line with experiments on E→D substitutions and the general similarity of force field parameters for D and E in the coarse-grained models used to generate the data (39, 41).

To explore how K→R substitutions compare with other residue types, we used ALBATROSS to predict *R*_*g*_ changes for all possible single-residue substitutions across sequences in the DisProt database (Fig. 2B). The results reveal that side-chain identity is not universally impactful. While the acidic D→E group remains in a “neutral” zone, substitutions involving basic, aromatic, or proline residues produce significantly broader distributions. This indicates that the specific chemistry of these side chains, such as the guanidinium group in arginine versus the amine in lysine, is a more potent regulator of IDP dimensions.

The K→R substitutions fall within this more sensitive landscape, providing statistical context for their moderate but systematic impact on the *α*Syn ensemble. It should be noted, however, that these broader trends are derived from a deep learning model trained on coarse-grained molecular dynamics simulations. Consequently, the observed differences reflect the underlying parameterization of amino acid force fields, representing a computational estimate of how side-chain chemistry influences polymer behavior.

### Electrostatic Screening Attenuates K→R Substitutions Compaction

To evaluate the role of electrostatic interactions in the compaction induced by K→R substitutions, we measured the *R*_*g*_ of the *α*Syn variants across increasing NaCl concentrations (50–300 mM) (Supplementary Table S2). In the absence of added salt, the wild-type (WT) protein exhibited an *R*_*g*_ of 29 Å, which is consistent with previous SAXS reports (64). This value progressively decreased to 26 Å for variant V7, consistent with increasing compaction upon K→R substitution (Fig. 1C). However, upon addition of 50 mM NaCl, all variants showed a clear expansion of the chain dimensions (Fig. 3A), with the WT increasing to 34 Å and V7 to 31 Å. The salt-induced expansion is consistent with the expected behavior of a polyampholyte polymer (65), and with previous SAXS experiments in 150 mM NaCl (66). The expansion further suggests that at low ionic strength, attractive intramolecular electrostatic interactions, particularly between oppositely charged domains, favor compaction. As a result, further increases in salt concentration continued to expand chain dimensions (Fig. 3A), reflecting the screening of electrostatic interactions. Despite this overall expansion, the compaction trend due to K→R substitutions was still clearly observed up to 150 mM added salt.

**Fig. 3.**
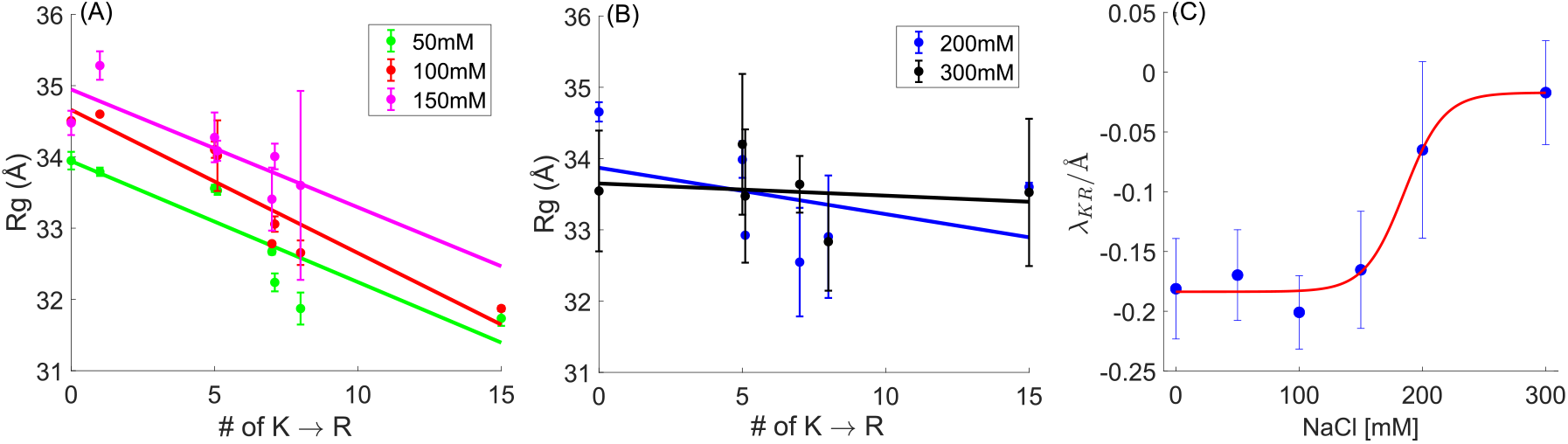
Lysine-to-arginine substitutions and ionic strength modulate *α*Syn compaction. (A) SAXS-derived radius of gyration (*R*_*g*_) for *α*Syn variants at 50–150 mM NaCl. *R*_*g*_ decreases with increasing K→R substitutions, with the degree of compaction modulated by ionic strength. (B) *R*_*g*_ of *α*Syn variants at 200 and 300 mM NaCl, showing smaller modulation of the K→R compaction trend at higher salt. (C) Dependence of the K→R compaction slope (*λ*_KR_) on salt concentration, highlighting the role of electrostatic screening in tuning *α*Syn conformational sensitivity. Errors are calculated as explained in the Methods section.

At 300 mM NaCl, the substitution-dependent trend was essentially abolished, with *R*_*g*_ values converging across variants (Fig. 3B). This suggests that the compaction induced by K→R substitutions depends on short-range electrostatic interactions, such as transient ionic bridges. At elevated salt concentrations, electrostatic interactions become strongly screened, reducing the formation of these short-range contacts and thereby diminishing the substitution-dependent differences in chain dimensions. Thus, the effect of K→R substitutions is most pronounced under conditions where local electrostatic interactions can occur, and diminishes under high-salt conditions where electrostatic interactions are strongly screened.

The salt dependency of K→R substitution effect shows a co-operative transition (Fig. 3C). Specifically, *λ*_*KR*_ as a function of added salt remains relatively constant at low salt concentrations and then decreases sharply between approximately 150 and 200 mM NaCl, approaching zero at higher ionic strength. Sigmoidal fitting captures this transition and indicates that the K→R dependent compaction is disrupted once electrostatic screening exceeds a critical threshold.

Comparison with the computational predictions from ALBATROSS revealed a systematic offset in absolute values, even under identical ionic strength conditions that it was trained on (150 mM NaCl). Specifically, ALBATROSS predicted *R*_*g*_ values ∼ 3 Å larger than those measured by SAXS (Fig. S4). Although ALBATROSS predictions correspond to 37 ^°^C, our standard SAXS measurements were performed at 20 ^°^C, additional SAXS measurements conducted at 37 ^°^C (0 and 50 mM NaCl) showed slightly smaller *R*_*g*_ values for all variants (Fig. S6), further increased the discrepancy with ALBATROSS predictions. However, the slope of the correlation (*λ*_*KR*_) was preserved, indicating that the relative scaling behavior across variants is captured despite the offset in magnitude.

### Modulation of the K→R Conformational Landscape

To gain deeper insight into how K→R substitutions in *α*Syn alter the conformational landscape, we applied molecular dynamics (MD) simulations, a diffusion model (STARLING) (45) and Ensemble Optimization Method (EOM) analyses (57), all developed specifically for IDPs. MD simulations, using the CALVADOS and Mpipi-Recharged coarse-grained force fields, revealed clear differences between the contact maps for WT and the V7 K→R variant (Fig. 4A-B, S7). In the absence of salt, contact maps showed a notable increase in long-range interactions for V7 compared with WT, indicating that K→R substitutions promote a more compact conformational ensemble. Addition of salt reduced, but did not fully eliminate, these contacts that are spaced far away along the protein chain, suggesting that electrostatic screening partially counteracts the compaction induced by arginine substitutions. For CALVADOS simulations, these observations persisted even after excluded-volume normalization of the contact maps (67) (Fig. S8), highlighting the changes in long-range contacts that arise from the interplay between increased residue ‘stickiness’ from arginine substitutions and increasing amounts of electrostatic screening.

**Fig. 4.**
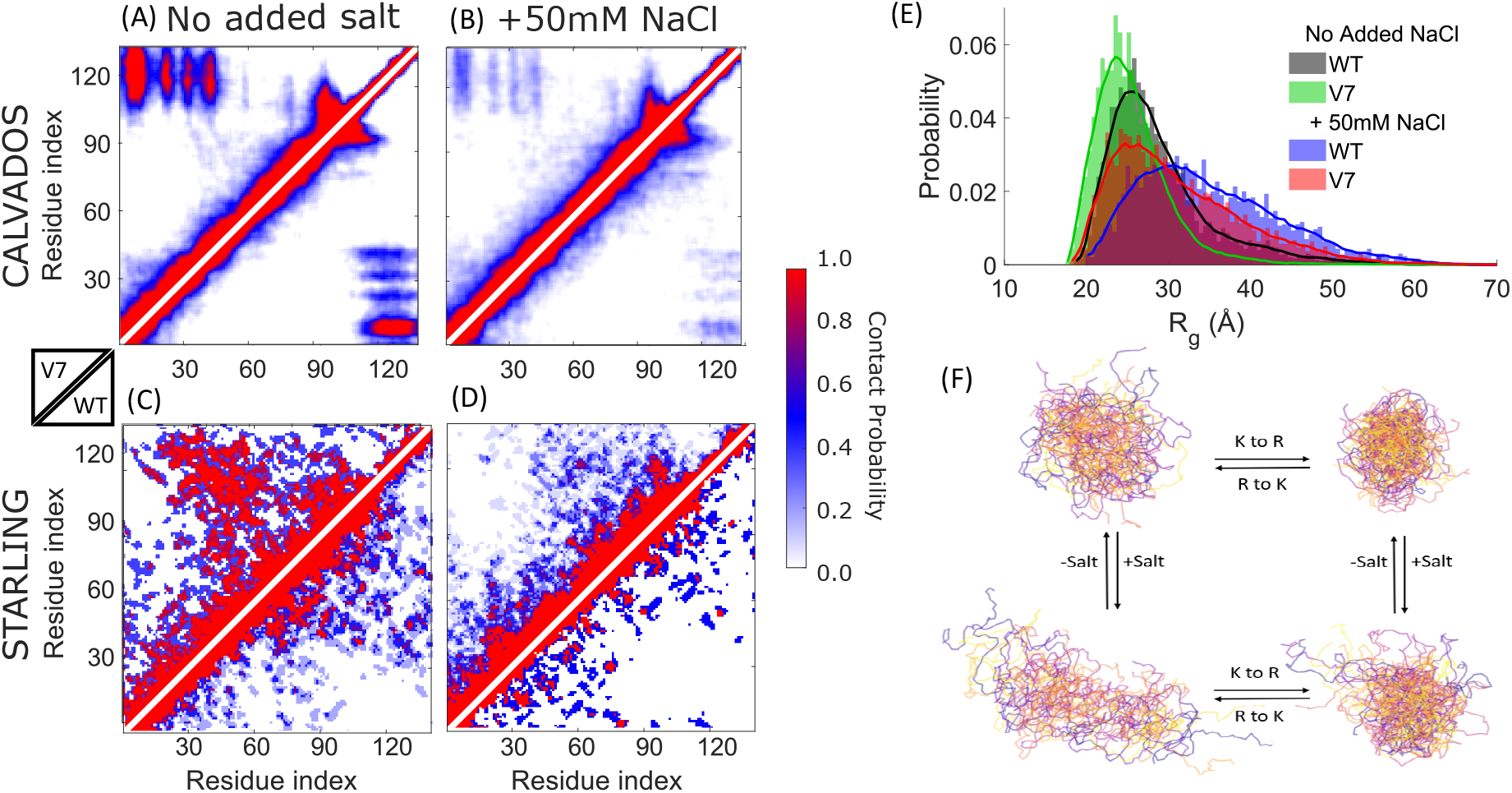
Salinity and K→R-dependent modulation of *α*Syn conformational ensemble. Contact maps from CALVADOS+BME simulations (A,B) and STARLING analysis (C,D) show *α*Syn without (A,C) and with 50 mM NaCl (B,D). In each map, the upper triangle portion represents V7 variant (all 15 lysine substituted to arginine), while the lower portion corresponds to WT. Contacts are normalized to the total number of contacts per ensemble. (E) *R*_*g*_ distributions from CALVADOS+BME analysis illustrate ensemble compaction for WT and V7 variant with and without 50mM NaCl. More pronounced differences between the distributions are observed at 150 mM NaCl (Fig. S9) (F) Representative structures from the MD simulation summarizing how K→R substitutions induce compaction, whereas increasing ionic strength expands the ensemble.

We then performed STARLING modeling that rapidly generated representative conformations of IDPs using a diffusion model trained by IDP coarse grained simulations (Fig. 4C-D). We calculated SAXS profiles from the structures generated using these tools, which were subsequently analyzed against our experimental data to post-select relevant represented frames (see Methods). This approach recapitulated the MD results, confirming that K→R substitutions favor a more compact ensemble, and that high ionic strength partially restores expansion.

Finally, we characterized the global dimensions of the protein by calculating the *R*_*g*_ distributions directly from the CALVADOS simulations weighted by Bayesian/Maximum Entropy (BME) to the SAXS data. This analysis provides a detailed view of ensemble heterogeneity, confirming that K→R substitutions shift the distribution toward more compact states (Figs. 4E, S9). To validate these computational results against our experimental data, we also applied the Ensemble Optimization Method (EOM), which selects sub-ensembles from a structural pool to fit the SAXS scattering profiles. The EOM analysis (Fig. S10) showed excellent agreement with the MD-derived distributions, maintaining the observed trends across all salinity range.

In all tested salt conditions, increasing the numbers of K→R substitutions led to narrower *R*_*g*_ distributions, indicating that arginine substitutions constrain the ensemble toward more uniform, compact conformations. This finding is consistent with the SAXS-derived average *R*_*g*_ values and supports the notion that arginine stabilizes specific conformational sub-populations.

However, a striking difference emerged when comparing the same variants between absent and added NaCl. The *R*_*g*_ distributions are consistently broader than at 0 mM NaCl for the same level of substitution. This suggests that increased salinity enhances ensemble heterogeneity, possibly by weakening long-range electrostatic interactions that otherwise help restrict the conformational space. Taken together with MD and STARLING analyses, these results highlight how K→R substitutions cooperatively shape the conformational landscape of *α*Syn across altered salinity (Fig. 4F).

### Arginine Substitution Alters *α*Syn Aggregation Propensity

The misfolding and aggregation of *α*Syn into amyloid fibrils is a hallmark of Parkinson’s disease and related synucleinopathies, driving neurodegeneration through the formation of toxic oligomers and insoluble deposits. Interestingly, nature has evolved *α*Syn as a protein entirely devoid of arginine residues, raising the question of how introducing arginine might influence its aggregation behavior. To investigate this, we performed Thioflavin T (ThT) fluorescence assays for all K→R variants. ThT fluorescence increases upon binding to amyloid fibrils, providing a reliable readout for *β*-sheet-rich aggregate formation.

Our results show that the rate of aggregation increased with the number of K→R substitutions (Fig. 5). Arginine-rich variants aggregated more rapidly, exhibiting shorter lag times and steeper growth phases. Notably, the fully substituted variant (V7) displayed the shortest lag time, indicating that K→R substitutions accelerate the initial nucleation events that drive *α*Syn aggregation.

**Fig. 5.**
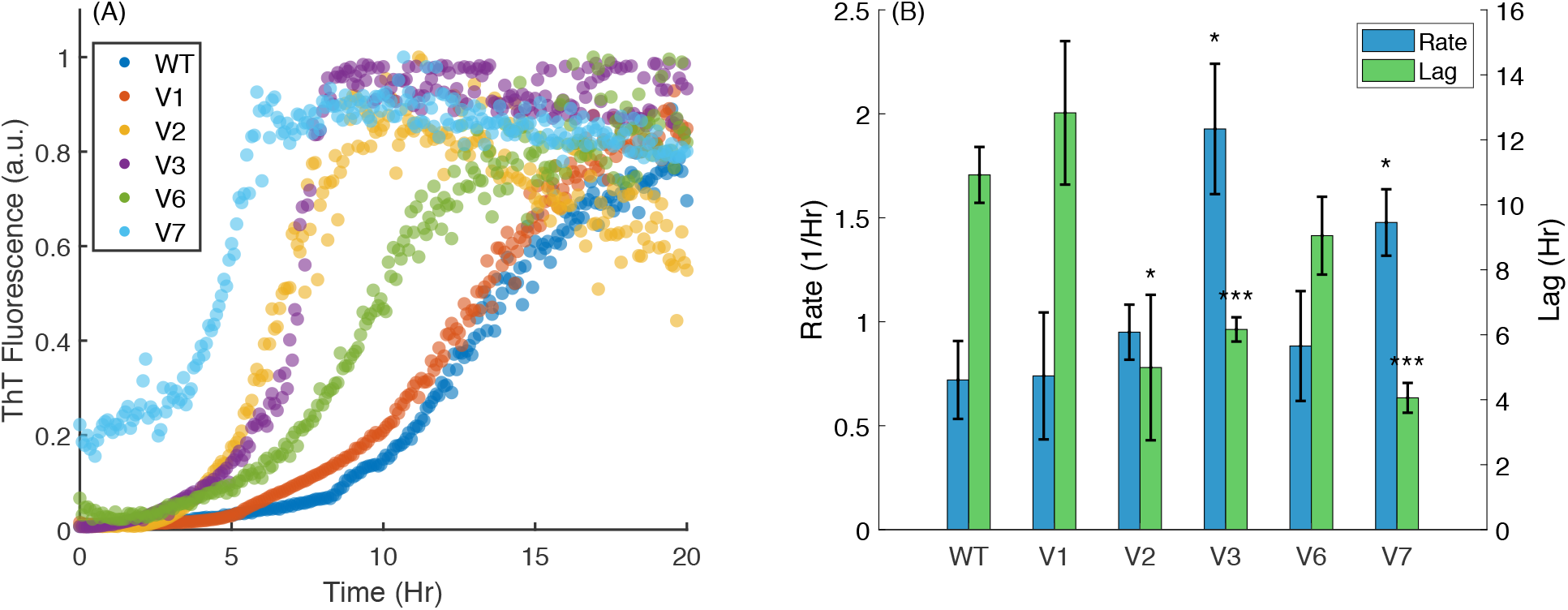
Aggregation assay of *α*Syn wild type (WT) and variants V1, V2, V3, V6 and V7. (A) Thioflavin T (ThT) fluorescence kinetics shows normalized aggregation curves for WT, *α*Syn variants over time. Data are averages of three repeats (37°C, 20 mM HEPES pH 7.4, orbital shaking). (B) Quantitative comparison of aggregation parameters derived from the ThT assay: aggregation rate (blue) and lag time (green). Error bars represent standard error across replicates. Statistical significance versus WT was assessed using two-sample t-tests and is indicated by asterisks (∗, ∗∗, ∗ ∗ ∗) corresponding to *p <* 0.05, *p <* 0.01, and *p <* 0.001, respectively.

## Discussion

Amino acid substitutions within a chemical group, such as lysine to arginine or glutamate to aspartate, are often considered interchangeable, yet their effects on conformational ensembles can differ markedly. In fact, using CALVADOS and ALBATROSS across the DisProt database we found a general feature of disordered sequence having structural sensitivity to the specific identity of basic residues, with the lysine-to-arginine ratio acting as a broad regulator of chain dimensions (Fig. 2A). Similar principles apply to hydrophobic residues, where small variations in side-chain size or branching can shift compaction and intermolecular interactions (Fig. 2B). These observations emphasize that not all substitutions within the same class behave equivalently, and systematic studies are needed to identify the most consequential alterations. Our experiments specifically address this challenge by focusing on these charge-preserving substitutions, which are among the most critical for understanding how the chemical identity of residues, beyond simple net charge, regulates IDP conformation and function.

IDPs, such as *α*Syn, exist as dynamic conformational ensembles whose dimensions are sensitive to both sequence composition and solvent environment, and the interplay between the two. In this study, we experimentally demonstrated that lysine-to-arginine (K→R) substitutions tune *α*Syn compaction and aggregation propensity, and that this tuning depends on the ionic strength. Although both residues are positively charged, their distinct physicochemical properties introduce substantial differences in intramolecular interactions. Our work adds to the growing body of evidence (21, 31, 32) that conformational control can be achieved through charge-preserving substitutions of charged residues, highlighting that the chemical “identity” serves to fine-tune the conformational ensemble beyond the dominant effects of net charge and patterning (68). This sensitivity to side-chain identity is not universal for all charged residues. For instance, previous studies on acidic residues have shown that glutamate-to-aspartate (E→D) substitutions result in no detectable change in the global conformational ensemble but only manifested itself in local structural changes (21).

The observed correlation between arginine content and *R*_*g*_ reduction likely stems from the unique properties of the guanidinium group. Unlike the primary amine of lysine, the planar guanidinium group of arginine facilitates cation-*π* interactions with aromatic residues (24) and attractive guanidinium– guanidinium interactions (32). The former is particularly relevant given the cluster of aromatic residues in the C-terminus (Y125, Y133, and Y136) which are known to engage in long-range contacts with the N-terminus (50, 69), and tyrosines are also the most conserved amino acids in the C-terminal domain (Fig. S12). By introducing arginines into the N-terminal lipid-binding domain, we likely enhance the affinity of this region for the C-terminal tyrosines, effectively “locking” the protein into more compact conformational states.

Beyond the importance of arginine content, our data hints at a positional effect of arginines; this is mostly clearly observed in two pairs of variants: V2/V3 (5 substitutions) and V4/V5 (7 substitutions). In both cases, the variants contain the same number of lysines and arginines but yield distinct conformational outcomes. Specifically, variants V2 and V4, which carry a higher density of substitutions within the N-terminal lipid-binding domain, exhibit lower *R*_*g*_ values than their counterparts, V3 and V5, where substitutions are distributed differently. Previous work has shown that it is possible to change compaction while preserving global amino acid composition via extensive permutation of the amino acid sequence (68, 70). Our work suggests that more subtle sequence variation and context can be used to modulate long-range interactions.

Additionally, the delocalized charge of the guanidinium group allows arginine to form stronger, more stable hydrogen bonds, enabling it to act as a hub for both local and non-local interactions. This is further compounded by the distinct hydration signatures of the two residues (20). Our SAXS and MD simulation data suggest that as K→R substitutions increase, the protein shifts from a highly hydrated, expanded coil toward a more collapsed ensemble, where water-mediated interactions are increasingly replaced by direct side-chain contacts.

The expansion of *α*Syn upon the addition of 50 mM NaCl confirms its polyampholytic behavior where attractive inter-domain charges are screened, consistent with recent small-angle neutron scattering observations (71). Remarkably, the K→R compaction effect (*λ*_*KR*_) persists up to 150 mM NaCl. This indicates that arginine-mediated compaction is not merely a product of long-range Debye-Hückel electrostatics, but involves short-range, high-affinity contacts (such as hydrogen bonding and cation-*π*) that are less sensitive to ionic strength. The MD and STARLING data suggest that these contacts are between residues that are spaced far away along the peptide chain (red markings in Fig. 4A-D). The “collapse” of the *λ*_*KR*_ parameter at 300 mM NaCl marks the critical threshold where even these robust short-range interactions are finally overwhelmed by the electrostatic screening.

To further contextualize these findings, we compared the *λ* parameters derived from our SAXS data against several state-of-the-art modeling approaches, which can be broadly categorized into two groups: de novo predictive simulations and experimentally-refined ensembles. The first group, which includes STARLING, ALBATROSS, and MD simulations with the CALVADOS or Mpipi force fields. In contrast, methods such as Guinier analysis, MFF, EOM, and the CALVADOS+BME framework incorporate SAXS experimental measurements to generate the ensemble characteristics (*R*_*g*_, *ν*) and relevant MD frames. While most models capture the general trend of *K* → *R* mediated compaction, there is a notable divergence in how they treat the sensitivity of these interactions to ionic strength (Fig. S13). Specifically, the experimental *λ* values show a distinct plateauing effect up to 150 mM NaCl before collapsing at 300 mM, a non-linear response that challenges models relying primarily on mean-field electrostatic approximations. Notably, our results demonstrate that CALVADOS, even without BME refinement to the SAXS data, captures the *K* → *R* trend at low salinity with high fidelity. This comparison highlights that while current force fields are increasingly adapt at modeling IDP dimensions, capturing the specific short-range “stickiness” of the guanidinium group in a saline environment remains a sensitive benchmark for model accuracy.

MD simulations, STARLING diffusion modeling, and EOM analysis provided ensemble-level insights beyond average *R*_*g*_ values. The molecular basis for these observations may be linked to the inherent stickiness of arginine residues. Indeed, certain hydropathy scales indicate that arginine possesses a stickiness value approximately 2.5-3.5 times greater than that of lysine (72, 73). This increased propensity for interaction likely drives the enhanced long-range contacts observed in our contact maps for the fully substituted variant (V7). Furthermore, moderate salt broadened our measured distributions, suggesting that electrostatic screening increases ensemble diversity by weakening the stabilizing intra-chain contacts provided by arginine.

Despite the well documented role of N- and C-terminal long-range interactions in shielding *α*Syn from aggregation (69, 74–77), arginine substitutions led to a paradoxical increase in the aggregation rate. This counterintuitive effect suggests that long-range interactions in *α*Syn do not inherently protect against fibrillization (51). Instead, arginine may stabilize specific compact sub-populations that increase the exposure or ‘nucleation-readiness’ of the NAC domain. This is consistent with recent MD simulations demonstrating that macromolecular crowding similarly shifts *α*Syn toward more compact conformations and can actually enhance aggregation propensity (78). Furthermore, as shown by our analysis, the restriction of ensemble heterogeneity likely reduces the entropic barrier required to reach the transition state for fibrillization, thereby accelerating the nucleation process. Additional analyses are, however, needed to understand which aggregation processes (54) are most strongly affected by the introduction of arginines.

Overall, our results demonstrate that chemically subtle substitutions between similarly charged residues can systematically reshape the energy landscape of IDPs. Nature’s total exclusion of arginine from the native *α*Syn sequence appears to be an evolutionary strategy to maintain ensemble breadth and solubility. Indeed, when we analysed 51 synuclein sequences from different species (79), we find on average 13 lysines per sequence but less than one arginine. By favoring lysine, the protein preserves a higher degree of conformational entropy, preventing the “trapping” of the protein in the compact, pro-aggregatory states observed in our arginine-rich variants. Furthermore, our approach of systematically exchanging residues with similar coarse-grained properties, yet distinct side-chains, provides a rigorous experimental benchmark for computational development. Such systematic datasets are essential for finetuning the next generation of MD force fields and AI-driven predictors, moving them beyond net-charge approximations toward a more nuanced, atomistic understanding of protein disorder. Ultimately, this work suggests that side-chain identity, in addition to global charge and patterning, is a critical, yet underappreciated, lever for modulating the dimensions and aggregation kinetics of IDPs.

## Materials and Methods

An extended description of protein purification, SAXS measurements, and additional data that contributes to the results is provided in SI Appendix. In brief, The recombinant proteins were produced in Escherichia coli and then purified by FPLC. SAXS measurements were performed at three different synchrotron facilities.

## ACKNOWLEDGMENTS

The synchrotron SAXS data were collected at beamline P12 operated by EMBL Hamburg at the PETRA III storage ring (DESY, Hamburg, Germany), at beamline B21 at Diamond Light Source (Didcot, UK), and at beamline BM29 at the European Synchrotron Radiation Facility (ESRF, Grenoble, France).This work benefited from access to the DESY,Hamburg, Germany, an Instruct-ERIC centre. Financial support was provided by Instruct-ERIC (PID 35548)

We would like to thank Katsuaki Inoue (Diamond Light Source), Hayden Fisher (ESRF), and Clément Blanchet, Cy Jeffries, and Dima Molodenskiy (EMBL) for their assistance in using the beamlines.

This work has been supported by the National Science Foundation under Grant No. MCB-2113302, the United States-Israel Bi-national Science Foundation under Grant No. 2020787, the Israel Science Foundation under Grants No. 1454/20. This work was supported by the PRISM (Protein Interactions and Stability in Medicine and Genomics) center funded by the Novo Nordisk Foundation (NNF18OC0033950 to K.L.-l.) and the support from Aufzien Family Center for the Prevention and Treatment of Parkinson’s Disease (APPD) at Tel Aviv University.

## Supporting Information for

### Methods

#### Protein purification

Protein purification followed periplasm purification of Huang et al.(1) with several modifications (2). The pT7-7 plasmid encoding *α*-Synuclein was a gift from Hilal Lashuel (Addgene plasmid # 36046) (3). Site-directed mutagenesis was carried out using PCRBIO HS Taq DNA Polymerase (PCR Biosystems Ltd.). Mutations were confirmed by sequencing the full-length protein-coding region of each mutant construct.

Chemically competent Escherichia coli BL21(DE3) cells were transformed with the modified vector and subsequently plated on agar plates containing 100*µg/ml* ampicillin. A single colony was picked to inoculate 50 ml LB medium (Difco) supplemented with 100*µg/ml* ampicillin and grown overnight at 37°C. The overnight culture was then used to inoculate 1 L LB medium containing 100*µg/ml* ampicillin in a baffled 5 L Erlenmeyer flask. Expression cultures were grown in a shaking incubator at 37°C at 220rpm for 2 hr until the optical density at 600 nm reached 0.7-1.0. Protein expression was induced by the addition of Isopropyl b-D-1-thiogalactopyranoside (IPTG) to a final concentration of 1 mM. The cultures were grown for 4 hr before harvesting.

Cells were pelleted by centrifugation and stored at -80°C for later use. For protein purification, cell pellet from a 4 L culture was resuspended in 200 ml of osmotic shock buffer (30 mM Tris–HCl pH 8.0, 30% sucrose, 2 mM EthyleneDiamineTetraacetic Acid (EDTA) and 5 mM 2-Mercaptoethanol) and incubated with gentle shaking for 20 min at 25°C. The suspension was centrifuged at 14,000 x g for 20 min at 4°C. Pallet was resuspended with 180 ml of ice-cold water containing 75 µL of saturated MgCl2, stirred on ice for 10 min, then treated with 10 mg/ml streptomycin sulfate for a additional 10 min stirring. The mixture was centrifuged for 10 min at 17,000 x g at 4°C and the supernatant was retained. A second centrifugation under the same condition for 30 min was preform to further clarify the solution. *α*-Synuclein protein was precipitated by salting-out with 54 gr ammonium sulfate and then dialyzed overnight against 20 mM Tris–HCl, pH 8.0 and 2 mM EDTA. After dialysis, the protein solution was stored at - 20 °C.

A final purification step was done using ion exchange chromatography. Protein sample was loaded onto a Q-Sepharose Fast Flow column (Amersham-Pharmacia Biotech) and eluted with a linear NaCl gradient from 0 to 0.5 M NaCl at 1 ml/min flow rate. The purified protein was stored at - 20 °C. Protein purity exceeded *>*90% as confirmed by SDS-PAGE (Fig. S11). Typical yields from 4 L cultures were 5–10 mg of pure protein.

#### Circular Dichroism

Far-UV circular dichroism spectra of *α*-Synuclein variants were measured at a concentration of 30 *µ*M in a 20 mM HEPES buffer pH = 7.2, 1mm cuvette at 20°C, using Chirascan Circular Dichroism Spectrometer (Applied Photophysics, Leatherhead, Surrey, UK). Each scan was recorded over the range of 190-260 nm, bandwidth of 1 nm, and an average time of 0.5 s per point. For each sample, three scans were measured and averaged.

#### SAXS measurements and analysis

All *α*-synuclein variants were dialyzed into a measurement buffer containing 20 mM HEPES (pH 7.2). Following dialysis, the dialysate was retained and utilized as the specific background buffer for subtraction to ensure precise solvent matching. Protein concentrations were determined using an extinction coefficient of 6000 M^−1^cm^−1^. Prior to measurements, samples were filtered using Amicon centrifugal filters (100 kDa cutoff) to remove potential higher-order aggregates. Measurement buffers were supplemented with 1 mM TCEP to minimize radiation-induced damage.

Final protein concentrations were adjusted to 0.5, 1, and 2 mg/mL. Scattering intensity curves were collected at these three concentrations to assess possible concentration dependent effects. All samples were measured at three synchrotron facilities: beamline B21 at Diamond Light Source (Didcot, UK), beamline P12 at EMBL at DESY (Hamburg, Germany), and beamline BM29 at the European Synchrotron Radiation Facility (Grenoble, France). Initial data processing involved the subtraction of the corresponding dialysis buffer from each protein scattering curve.

The radius of gyration (*R*_*g*_) for each concentration was determined via Guinier analysis (4) by performing a linear regression of ln[*I*(*q*)] versus *q*^2^ in the low-*q* region (*q* · *R*_*g*_ *<* 1.3). The uncertainty in 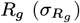 was propagated from the standard error of the slope (*σ*_*slope*_) as follows:

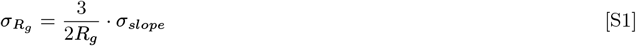

To account for interparticle effects and ensure optimal signal-to-noise, final structural parameters were obtained by linear extrapolation to a concentration of 1 mg/mL. A weighted linear least-squares fit was employed, where weights were defined as the inverse variance of each measurement 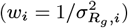. The uncertainty of the extrapolated value (*σ*_*extrap*_) was calculated using the full covariance matrix of the fit to account for the correlation between the intercept (*b*) and the slope (*m*):

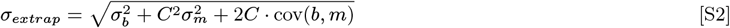

To quantify the structural impact of Lysine-to-Arginine (K→R) substitutions across the *α*-synuclein variants, the extrapolated *R*_*g*_ values were modeled as a function of the number of substitutions (*N*_*K*→*R*_). A secondary linear regression was performed:

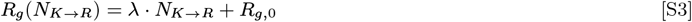

where *λ* represents the rate of change in the radius of gyration per substitution, and *R*_*g*,0_ is the intercept representing the *R*_*g*_ of the wild-type or reference variant.

The final sensitivity parameter, *λ*, was determined by a weighted linear regression of the extrapolated *R*_*g*_ values against the number of K→R substitutions. To ensure that variants with higher experimental precision contributed proportionally to the model, weights were defined as the inverse variance 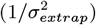 of the extrapolated *R*_*g*_ values. The uncertainty *σ*_*λ*_ was reported as the standard error of the weighted slope estimate.

#### CALVADOS *α*Syn simulations

Molecular dynamics simulations were performed using OpenMM (version 8.2.0) (5). Simulations were conducted in the NVT ensemble using a Langevin integrator with a time step of 10 fs and friction coefficient of 0.01 ps^−1^. The *α*Syn variants were simulated in a cubic box with an edge length of 56.82 nm under periodic boundary conditions. Simulations were performed at a temperature of 293.15 K (20^°^C), pH 7.5 and varying levels of ionic strength (including the contribution from experimental buffer salinity): 10 mM, 60 mM, 100 mM, 150 mM, 200 mM, 300 mM.

Each system was simulated in 3 replicas for 7.07 *×* 10^6^ simulation steps, corresponding to 70.7 ns of simulated time. Trajectories were written to DCD files every 7 *×* 10^3^ steps, resulting in 1010 frames per replica. Final simulation trajectories for each *α*Syn variant at each salt concentration contained 3000 weakly correlated frames, after discarding the initial 10 frames in each replica and concatenating the rest.

We used the CALVADOS 3 coarse-grained model (6), where each amino acid residue is mapped onto a single bead and solvent particles are implicitly considered.

Bonded interactions are modelled using a harmonic potential:

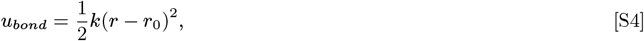

with force constant *k* = 8033 kJ mol^−1^ nm^−2^, and equilibrium distance *r*_0_ = 0.38 nm.

Non-bonded non-ionic interactions are modelled using an Ashbaugh-Hatch (AH) potential (7, 8), truncated and shifted at *r*_*c*,AH_ = 2 nm:

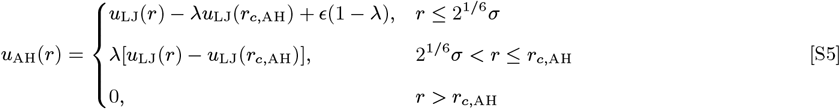

Here, *σ* = (*σ*_*i*_ + *σ*_*j*_)*/*2 and *λ* = (*λ*_*i*_ + *λ*_*j*_)*/*2 for residues *i* and *j*. The residue-specific diameters (*σ*_*i*_, *σ*_*j*_) and stickiness values (*λ*_*i*_, *λ*_*j*_) are key parameters that influence the strength and range of AH interactions. *u*_LJ_ is the classic Lennard-Jones (LJ) potential:

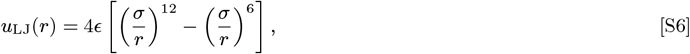

where *ϵ* = 0.8368 kJ mol^−1^.

Salt-screened electrostatic interactions are modelled using a Debye-Hückel potential, truncated and shifted at *r*_*c*,DH_ = 4 nm:

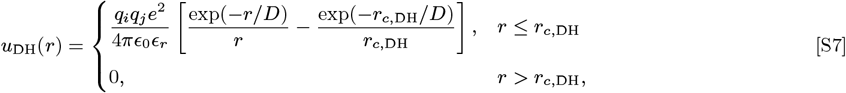

where *e* is the elementary charge, *q*_*i*_ and *q*_*j*_ are the charges of residues *i* and *j, ϵ*_0_ is the vacuum permittivity, 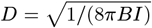 is the Debye length of an electrolyte solution of ionic strength *I*, and *B*(*ϵ*_*r*_) is the Bjerrum length of the temperature-dependent dielectric constant of the implicit aqueous solution, *ϵ*_*r*_, modelled by the empirical relationship (9):

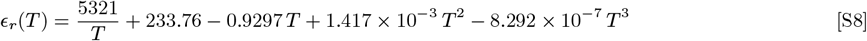

The partial charge of Histidine residues is set according to the simulated pH, using the Henderson-Hasselbalch equation:

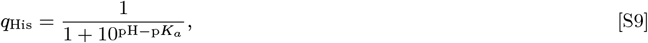

with *pK*_*a*_ = 6.0 (10).

#### CALVADOS Excluded-Volume Simulations

To quantify the amount of “residual structure” relative to a reference state, we generated excluded-volume (EV) models (11)for the WT and V7 *α*Syn variants using simulations with a modified version of CALVADOS 3 that models non-bonded interactions using only the repulsive portion of the Ashbaugh-Hatch potential. In practice, we set *q* = 0 and *λ* = 0 for all residue types, effectively removing the electrostatics contribution from the Debye-Hückel potential and the attractive portion of the Ashbaugh-Hatch potential, respectively.

EV simulations were conducted in the NVT ensemble using a Langevin integrator with a time step of 10 fs and friction coefficient of 0.01 ps^−1^. Single-chain *α*-synuclein was simulated in a cubic box with an edge length of 56.82 nm under periodic boundary conditions. Simulations were performed at a temperature of 293.15 K (20^°^C), pH 7.5, 0 M ionic strength, and with both termini left uncharged. Each system was simulated in 25 replicas for 140.07 *×* 10^6^ simulation steps, corresponding to 1400.7 ns of simulated time. Trajectories were written to DCD files every 7 *×* 10^3^ steps, resulting in 20010 frames per replica. The final EV simulation trajectories for WT and V7 *α*Syn each contained 500,000 weakly correlated frames, after discarding the initial 10 frames in each replica and concatenating the rest, totalling ∼ 35 *µ*s of simulated time.

#### Contact Map Excluded-Volume Normalization

We calculated inter-residue contact frequency maps from the CALVADOS EV simulation trajectories and used them to normalize the values observed from the CALVADOS 3 simulations (11):

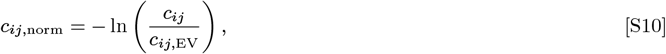

where *c*_*ij*_ is the contact frequency for residue pair *ij* in the CALVADOS 3 simulation and *c*_*ij*,EV_ is the contact frequency for the same residue pair in the CALVADOS EV simulation.

#### Bayesian/Maximum Entropy (BME) Reweighting

We first converted the CALVADOS coarse-grained simulation trajectories to all-atom using cg2all (12) (*CalphaBasedModel*) and then back-calculated SAXS profiles from the all-atom trajectories using Pepsi-SAXS (13) with default parameters. Simulated trajectories were then reweighted following the Bayesian/Maximum Entropy (BME) reweighting approach (14), aiming to maximize the agreement between calculated and experimental SAXS observables while minimally changing the original simulation trajectory. This can be interpreted as a minimization problem: we change the initial set of uniform frame weights *w*^0^ towards a set of optimal weights (*w*_1_ … *w*_*n*_) that minimize function ℒ:

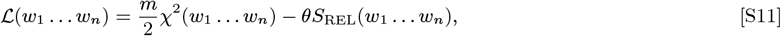

where *m* is the number of experimental measurements, *n* is the number of trajectory frames and *θ* is a global scaling parameter.

The relative entropy term, *S*_REL_, measures the deviation from the initial set of weights:

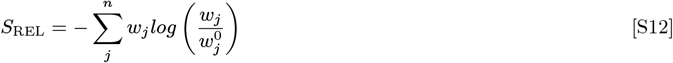

The decrease in uniformity of the weights corresponds to a decrease in the number of trajectory frames that effectively contribute to the calculated ensemble averages, and can be quantified by the fraction of effective frames, *ϕ*_eff_:

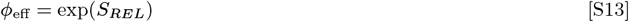

The level of experimental agreement is quantified by the reduced *χ*^2^:

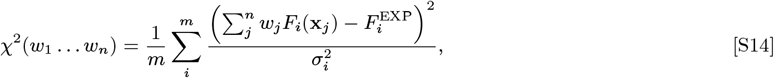

where 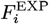 is the experimental measurement, *F*_*i*_ (x_*j*_) is the theoretical measurement calculated from each conformer, and 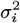 is the uncertainty associated with 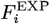, including experimental errors and inaccuracies introduced by the calculation of *F*_*i*_ (x_*j*_). The global scaling parameter *θ* is introduced because 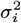 is not always known accurately (14). When *θ* →∞ we have no confidence in the experimental data. On the other hand, when *θ* → 0 a perfect match with experimental data is achieved.

The global scaling parameter *θ* must be adequately tuned so that the reweighted trajectory is in good agreement with the experimental data (low *χ*^2^) while minimally perturbing the initial set of frame weights (high *ϕ*_eff_). In practice, we can plot *χ*^2^ vs. *ϕ*_eff_ across different *θ* values, and look for the optimal *θ*, i.e. the one corresponding to the point of maximum curvature of the resulting curve, where we cannot optimize experimental agreement further without risking overfitting to the experimental data.

#### CALVADOS DisProt simulations

We first simulated each native (WT) IDR sequence in the DisProt database. For each WT sequence, two additional variants were simulated: (1) a lysine-to-arginine (K→R) substitution variant in which all lysine residues were replaced by arginine, and (2) an arginine-to-lysine (R→K) substitution variant in which all arginine residues were replaced by lysine. Thus, three sequences (WT, K→R, and R→K) were simulated per DisProt IDR entry.

Unless explicitly stated otherwise, we followed the same scheme as for the *α*Syn simulations. Each sequence was simulated in 1 replica. Each sequence composed of *N* residues was simulated in a cubic box with edge length *L* = (*N* − 1) *×* 0.38 + 4 nm under periodic boundary conditions. After applying this formula, when *L <* 8 nm, *L* was set to 12 nm. Conversely, when *L >* 900 nm, *L* was set to 900 nm. Sequences with *N >* 150 were simulated for ∼ 303 *× N* ^2^ simulation steps, and conformations were written to a DCD file every ∼ 0.3 *× N* ^2^ simulation steps. Final simulation trajectories for each variant of each IDR in the DisProt database contained 1000 weakly correlated frames, after discarding the initial 10 frames.

#### Ensemble optimization method (EOM)

In the ensemble optimization method (EOM), a pool containing 10,000 possible conformations based on protein sequence is generated with RANCH program (version 2.1) followed by scattering amplitudes calculation (15). Then, the ensemble (sub-group of pool conformations) whose combined theoretical scattering intensity best describes the experimental SAXS data is selected with a genetic algorithm (GAJOE program version 2.1) (15).

#### Thioflavin T (ThT) Aggregation assay

Aggregation was monitored over 20 hours at 37°C with the addition of 3 mm glass beads to improve reproducibility (16). Reactions were performed in 0.2 M HEPES buffer at a protein concentration of 1.5 mg/mL with 20 *µ*M Thioflavin T (ThT). Fluorescence measurements were performed using monochromators for both excitation and emission on a TECAN Infinite M200PRO plate reader. ThT was excited at 430 nm with a 20 nm bandwidth, and emission was recorded at 480 nm with a 20 nm bandwidth. Measurements were performed in triplicate, and ThT fluorescence intensity was recorded at 5 min intervals to capture the aggregation dynamics across lag, elongation, and plateau phases. The time-dependent fluorescence signal was fitted to a sigmoidal growth model using nonlinear least-squares regression. The fitting function was defined as:

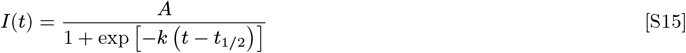

where *I*(*t*) is the ThT fluorescence intensity at time *t, A* is the maximal fluorescence intensity (plateau value), *k* is the apparent growth rate constant, and *t*_1*/*2_ is the half-time of aggregation (defined as the time at which the fluorescence reaches 50% of *A*).Each condition was analyzed in triplicate within each experimental repeat, and the resulting kinetic parameters were collected for subsequent statistical analysis.

#### ALBATROSS-Based Sequence-Wide Substitution Analysis

All proteins with disordered region (IDR) sequences were obtained from DisProt database. Only annotated disordered regions were used for subsequent analysis. For each native (WT) sequence, two additional variants were computationally generated: (1) a lysine-to-arginine (K→R) substitution variant in which all lysine residues were replaced by arginine, and (2) an arginine-to-lysine (R→K) substitution variant in which all arginine residues were replaced by lysine. Thus, three sequences (WT, K→R, and R→K) were analyzed per protein.

Sequence-based conformational predictions were performed using ALBATROSS with default parameters. For each of the three sequences per protein, the predicted radius of gyration (*R*_*g*_) was extracted. A linear fit was then applied to the three *R*_*g*_ values to determine a sequence-specific scaling coefficient (*λ*), reflecting the sensitivity of chain dimensions to the K/R compositional shift. This procedure was systematically repeated for all possible single–amino acid substitutions (*i*.*e*., replacing all residues of a given amino acid type with another type), generating analogous triplets of sequences per protein and extracting *λ* values for each substitution pair.

For each amino acid replacement category, *λ* values were aggregated across all proteins in the dataset. The distribution of *λ* values was analyzed by calculating the mean and standard deviation. Finally, a two-dimensional map was constructed in which each axis corresponds to the original and substituted amino acid type, and each cell represents the mean *λ* value across the full protein dataset (Fig. 2B).

#### STARLING Ensemble Generation and SAXS-Based Selection

Conformational ensembles were generated using STARLING for the WT sequence and all sequence variants. For each construct, 5,000 conformations were sampled. Residue–residue contact maps were calculated for every generated structure, where a contact was defined as a pair of residues with a *C*_*α*_–*C*_*α*_ distance shorter than 8 Å. Each individual conformation was subsequently converted into a theoretical SAXS profile using the EOM framework. This procedure resulted in 5,000 theoretical scattering curves per construct. Following the theoretical SAXS profiles were compared to the corresponding experimental SAXS data by *χ*^2^ minimization. Approximately 500 conformations yielding the lowest *χ*^2^ values were selected.

Finally, the contact maps corresponding to these ∼ 500 best-fitting conformations were averaged to generate the ensemble-averaged contact map presented in Figure 4. This approach ensures that the reported contact map represents conformations consistent with the experimental SAXS data.

#### Mpipi-recharged simulations

All simulations were performed using the OpenMM molecular simulation engine (version 8.0.0.dev-a780005)(5) through the OpenMpipi interface (version 0.1.0), running under Python 3.9.20. Simulations were executed on NVIDIA A100-SXM4-80GB GPUs using the CUDA platform (CUDA 12.2) with mixed precision.

Protein models were constructed using the OpenMpipi IDP builder, which generates a coarse-grained representation of the protein directly from its amino-acid sequence and initializes the system in a random-coil conformation. Systems were simulated using the *Mpipi-Recharged* coarse-grained force field, a refined variant of the Mpipi model that improves the description of electrostatic interactions in intrinsically disordered proteins (17).

Simulations were carried out under implicit solvent conditions with periodic boundary conditions enabled and a specified salt concentration for each system. Electrostatic interactions were modeled using Debye–Hückel screening according to the Mpipi-Recharged parameterization, which accounts for ionic strength in the solution. Center-of-mass motion removal was enabled during the simulations.

The equations of motion were integrated using the Langevin middle integrator with a time step of 10 fs. Temperature was maintained at 293.15 K (20^°^C) using a Langevin thermostat with a friction coefficient of 0.01 ps^−1^. Prior to production dynamics, each system underwent energy minimization to remove unfavorable contacts from the initial random configuration. Production simulations were performed in the NVT ensemble for 5.0 *×* 10^6^ integration steps, corresponding to approximately 50 ns of simulated time. Trajectories were written every 10 000 steps to DCD files for subsequent analysis. Thermodynamic quantities, including simulation step, time, potential energy, and instantaneous temperature, were recorded at the same interval using the openmm `StateDataReporter`.

To calculate the scaling exponent *ν* from the Molecular Dynamics trajectories, we analyzed the relationship between the average internal distance *R*(*s*) and the residue separation *s* = |*i* − *j*|. For each protein variant and salt concentration, the distance map was used to compute the mean distance for each separation *s* by averaging the *s*-th super-diagonal of the matrix, such that 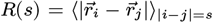. The scaling exponent was then extracted by fitting the data to the power-law relation 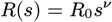 via linear regression in log-log space. To ensure the robustness of the fit and account for potential sensitivity to the fitting range, *ν* was calculated over five overlapping windows: [15, 40], [20, 40], [20, 45], [15, 50], and [25, 45]. The final values are reported as the mean *ν* across these ranges, with the uncertainty defined by the standard deviation *σ*_*ν*_.

## Supporting information

**Fig. S1.**
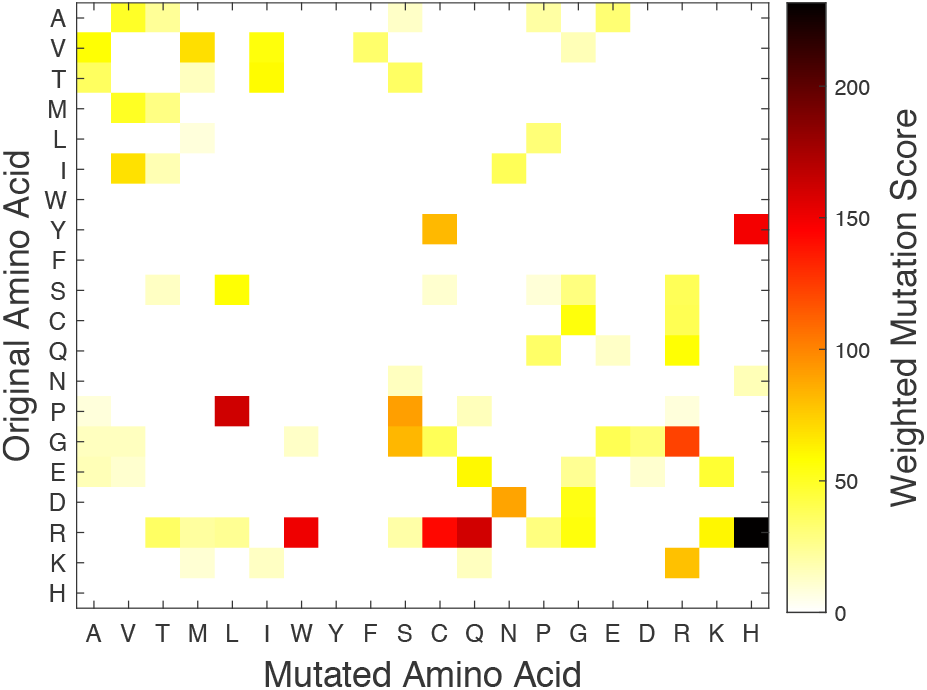
Distribution of disease-associated amino acid substitution frequencies within intrinsically disordered regions (IDRs). The analysis includes reported sequence variants and annotated disease-associated mutations curated in UniProt. The heatmap illustrates the frequency of single-residue substitutions for all proteins in the UniProt database within disordered regions. The vertical axis represents the original amino acid residue, while the horizontal axis represents the mutated residue. Each cell represents the total count of a specific mutation normalized by the abundance of the original amino acid within its corresponding sequence. The normalized score was calculated as the total number of observed substitutions from amino acid i to j divided by the fractional abundance of amino acid i (defined as the number of residues i divided by the sequence length).

**Fig. S2.**
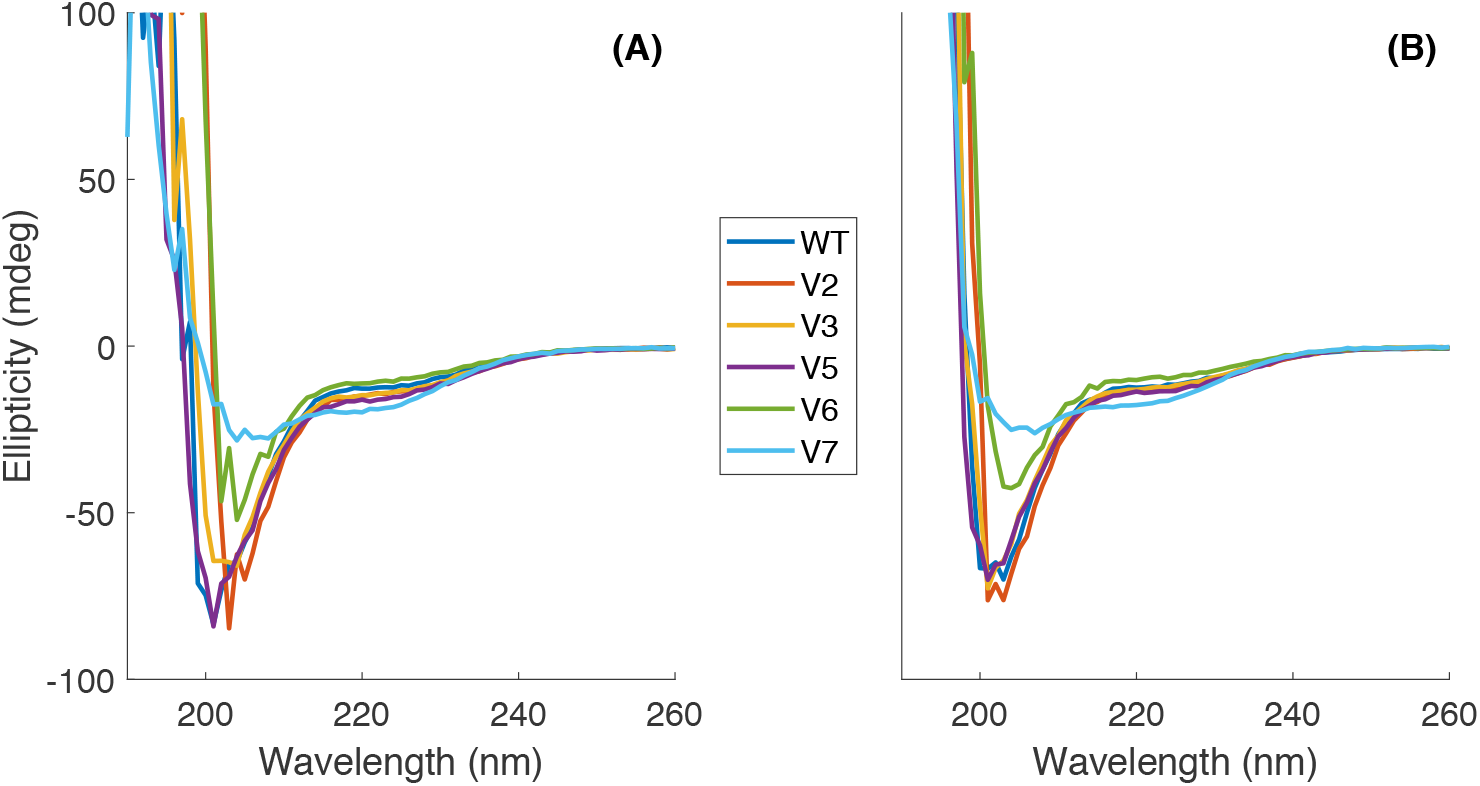
Circular dichroism (CD) spectra of *α*-synuclein (WT) and its variants recorded in 5 mM NaPO_4_ buffer (pH 7.2). (A) Spectra measured in the absence of NaCl (0 mM). (B) Spectra measured in the presence of 150 mM NaCl. All proteins exhibit a characteristic random coil spectrum, indicating the absence of stable secondary structure under both conditions. Secondary structure estimation was performed using BestSel (18).

**Table S1.**
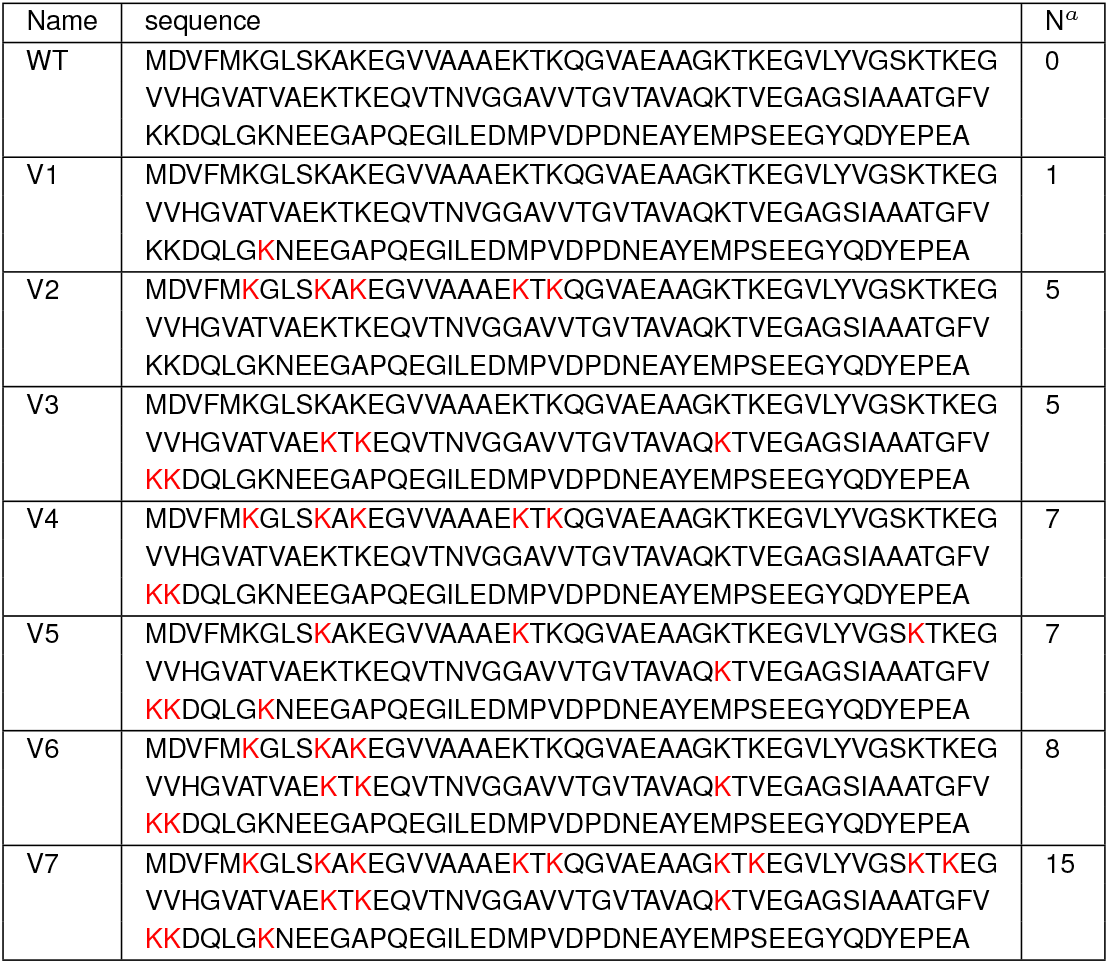
*α*-synuclein variants sequence used for SAXS measurements. *α*-synuclein sequence is drived from UniProtKB P37840. Marked in red are the Lysine-to-Arginine substitutions. *N*^*a*^ is the number of Lysine-to-Arginine substitutions sites.

**Fig. S3.**
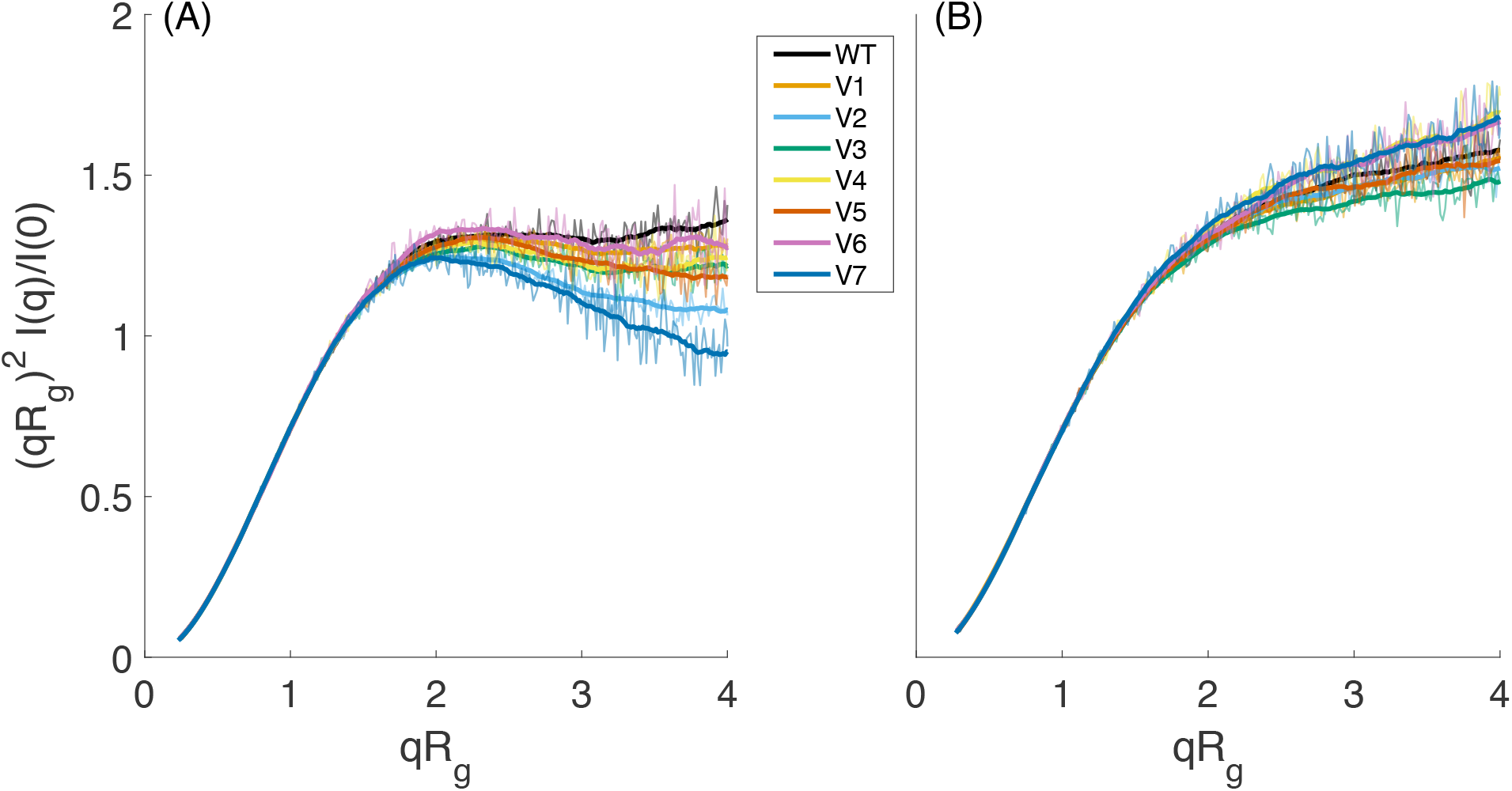
Kratky plots of *α*-synuclein (WT and variants). (A) 0 mM NaCl. (B) 150 mM NaCl. All proteins display a monotonic increase at high q values without a defined peak, consistent with the characteristic structural signature of an intrinsically disordered protein.

**Table S2.** Summary of *R*_*g*_ for *α*-synuclein variants across experimental and computational methods at 0-300mM NaCl. All values are reported in Å. Experimental *R*_*g*_ values were derived from SAXS Guinier analysis, while computational values were obtained from ALBATROSS predictions, STARLING modeling, Ensemble Optimization Method (EOM) analysis, and Molecular Dynamics (MD) simulations.

| 0 mM NaCl |  |  |  |  |  |  |  |  |  |
| --- | --- | --- | --- | --- | --- | --- | --- | --- | --- |
| Name | $R_g$ SAXS | $R_g$ CALVADOS | $R_g$ CALVADOS-SAXS | $R_g$ EOM | $R_g$ STARLING | $R_g$ Mpipi | $\nu$ Mpipi | $R_g$ MFF | $\nu$ MFF |
| WT | 29.37±0.48 | 28.86±0.69 | 30.21±0.70 | 30.77 | 36.25 | 33.63 | 0.49 | 31.85 | 0.53 |
| V1 | 29.57±0.52 | 29.18±0.82 | 30.10±0.83 | 31.03 | 36.33 | 33.96 | 0.48 | 32.54 | 0.48 |
| V2 | 27.89±0.20 | 27.53±0.88 | 28.35±0.89 | 28.91 | 35.65 | 28.47 | 0.42 | 29.93 | 0.42 |
| V3 | 28.56±0.89 | 28.25±0.75 | 28.54±0.78 | 29.06 | 35.70 | 31.75 | 0.43 | 32.37 | 0.46 |
| V4 | 28.01±0.03 | 26.81±0.84 | 28.43±0.83 | 29.20 | 35.04 | 25.85 | 0.38 | 30.60 | 0.47 |
| V5 | 28.77±0.07 | 27.95±0.81 | 29.64±0.81 | 30.30 | 35.63 | 28.15 | 0.39 | 30.78 | 0.41 |
| V6 | 27.07±0.14 | 26.89±0.87 | 28.77±0.89 | 29.55 | 35.31 | 26.48 | 0.35 | 29.73 | 0.49 |
| V7 | 26.55±0.15 | 25.48±1.11 | 26.70±1.12 | 27.59 | 33.86 | 23.36 | 0.31 | 28.39 | 0.39 |
| 50 mM NaCl |  |  |  |  |  |  |  |  |  |
| Name | $R_g$ SAXS | $R_g$ CALVADOS | $R_g$ CALVADOS-SAXS | $R_g$ EOM | $R_g$ STARLING | $R_g$ Mpipi | $\nu$ Mpipi | $R_g$ MFF | $\nu$ MFF |
| WT | 33.95±0.12 | 35.05±0.56 | 35.77±0.61 | 36.36 | 36.28 | 39.48 | 0.55 | 37.60 | 0.52 |
| V1 | 33.79±0.09 | 34.76±0.65 | 35.34±0.68 | 36.69 | 36.25 | 37.41 | 0.54 | 37.04 | 0.53 |
| V2 | 33.56±0.05 | 32.70±0.66 | 35.90±0.67 | 37.31 | 35.62 | 35.60 | 0.52 | 36.91 | 0.52 |
| V3 | 33.53±0.06 | 33.95±0.62 | 35.49±0.64 | 35.71 | 35.71 | 35.65 | 0.49 | 36.27 | 0.53 |
| V4 | 32.67±0.04 | 33.13±0.67 | 33.51±0.75 | 34.53 | 34.98 | 34.90 | 0.51 | 34.00 | 0.52 |
| V5 | 32.24±0.13 | 32.79±0.66 | 35.41±0.67 | 35.94 | 35.70 | 35.77 | 0.51 | 35.51 | 0.54 |
| V6 | 31.87±0.22 | 33.20±0.64 | 34.40±0.66 | 35.09 | 35.49 | 34.14 | 0.48 | 34.60 | 0.49 |
| V7 | 31.74±0.11 | 31.12±0.78 | 32.93±0.80 | 33.78 | 33.96 | 27.50 | 0.39 | 33.59 | 0.48 |
| 100 mM NaCl |  |  |  |  |  |  |  |  |  |
| Name | $R_g$ SAXS | $R_g$ CALVADOS | $R_g$ CALVADOS-SAXS | $R_g$ EOM | $R_g$ STARLING | $R_g$ Mpipi | $\nu$ Mpipi | $R_g$ MFF | $\nu$ MFF |
| WT | 34.51±0.02 | 36.33±0.58 | 37.70±0.60 | 37.43 | 37.19 | 40.52 | 0.58 | 36.79 | 0.60 |
| V1 | 34.60±0.00 | 35.73±0.61 | 37.39±0.62 | 38.12 | 36.81 | 40.22 | 0.57 | 38.38 | 0.57 |
| V2 | 34.10±0.11 | 35.07±0.70 | 36.64±0.73 | 37.33 | 36.44 | 37.67 | 0.54 | 37.03 | 0.60 |
| V3 | 34.08±0.50 | 35.01±0.63 | 35.46±0.64 | 36.07 | 36.35 | 37.97 | 0.53 | 37.87 | 0.52 |
| V4 | 32.78±0.02 | 34.43±0.64 | 35.09±0.66 | 36.85 | 36.06 | 37.80 | 0.55 | 34.56 | 0.60 |
| V5 | 33.06±0.11 | 35.04±0.68 | 35.55±0.71 | 36.41 | 36.45 | 37.48 | 0.54 | 34.37 | 0.60 |
| V6 | 32.66±0.17 | 34.25±0.68 | 35.02±0.69 | 35.72 | 35.25 | 37.95 | 0.54 | 34.92 | 0.60 |
| V7 | 31.87±0.03 | 33.30±0.59 | 34.13±0.61 | 34.82 | 34.90 | 32.66 | 0.48 | 32.68 | 0.60 |
| 150 mM NaCl |  |  |  |  |  |  |  |  |  |
| Name | $R_g$ SAXS | $R_g$ CALVADOS | $R_g$ CALVADOS-SAXS | $R_g$ EOM | $R_g$ STARLING | $R_g$ Mpipi | $\nu$ Mpipi | $R_g$ MFF | $\nu$ MFF |
| WT | 34.48±0.17 | 36.92±0.53 | 37.41±0.58 | 37.92 | 37.24 | 40.29 | 0.58 | 37.88 | 0.57 |
| V1 | 35.28±0.20 | 36.39±0.63 | 38.02±0.64 | 37.76 | 37.00 | 40.89 | 0.60 | 38.02 | 0.62 |
| V2 | 34.27±0.35 | 35.23±0.65 | 36.90±0.68 | 37.67 | 36.57 | 39.17 | 0.57 | 37.29 | 0.59 |
| V3 | 34.09±0.14 | 35.51±0.53 | 36.18±0.60 | 36.91 | 36.57 | 38.03 | 0.54 | 37.53 | 0.55 |
| V4 | 33.41±0.44 | 35.41±0.67 | 35.45±0.72 | 36.97 | 36.15 | 37.71 | 0.55 | 35.23 | 0.58 |
| V5 | 34.01±0.18 | 35.44±0.58 | 36.04±0.62 | 36.55 | 36.25 | 38.57 | 0.55 | 37.46 | 0.56 |
| V6 | 33.60±1.33 | 35.64±0.57 | 37.30±0.57 | 37.84 | 36.21 | 37.97 | 0.55 | 33.94 | 0.58 |
| V7 |  | 33.67±0.65 |  |  | 35.07 | 34.97 | 0.52 |  |  |
| 200 mM NaCl |  |  |  |  |  |  |  |  |  |
| Name | $R_g$ SAXS | $R_g$ CALVADOS | $R_g$ CALVADOS-SAXS | $R_g$ EOM | $R_g$ STARLING | $R_g$ Mpipi | $\nu$ Mpipi | $R_g$ MFF | $\nu$ MFF |
| WT | 34.66±0.14 | 37.34±0.65 | 35.74±0.68 | 35.18 | 37.34 | 40.19 | 0.58 | 36.55 | 0.61 |
| V1 |  | 37.09±0.61 |  |  | 37.14 | 40.81 | 0.59 |  |  |
| V2 | 33.99±0.26 | 36.08±0.54 | 37.05±0.58 | 35.89 | 36.65 | 39.79 | 0.58 | 40.58 | 0.45 |
| V3 | 32.92±0.00 | 35.87±0.64 | 35.91±0.71 | 35.47 | 36.65 | 39.40 | 0.57 | 39.31 | 0.39 |
| V4 |  | 36.06±0.66 |  |  | 36.46 | 39.41 | 0.58 |  |  |
| V5 | 32.55±0.76 | 35.25±0.57 | 36.31±0.62 | 33.30 | 36.16 | 39.01 | 0.57 | 34.74 | 0.58 |
| V6 | 32.90±0.86 | 34.98±0.69 | 33.15±0.72 | 34.21 | 36.18 | 38.54 | 0.57 | 34.32 | 0.63 |
| V7 | 33.60±0.06 | 34.22±0.63 | 34.12±0.68 | 34.43 | 35.14 | 36.33 | 0.55 | 37.22 | 0.53 |
| 300 mM NaCl |  |  |  |  |  |  |  |  |  |
| Name | $R_g$ SAXS | $R_g$ CALVADOS | $R_g$ CALVADOS-SAXS | $R_g$ EOM | $R_g$ STARLING | $R_g$ Mpipi | $\nu$ Mpipi | $R_g$ MFF | $\nu$ MFF |
| WT | 33.55±0.85 | 37.07±0.60 | 35.97±0.63 | 35.51 | 37.31 | 41.14 |  | 36.24 | 0.66 |
| V1 |  | 37.25±0.66 |  |  | 37.14 | 40.15 |  |  |  |
| V2 | 34.20±0.99 | 36.17±0.57 | 37.72±0.63 | 38.63 | 36.75 | 39.94 |  | 41.76 | 0.39 |
| V3 | 33.47±0.40 | 36.05±0.72 | 35.63±0.78 | 32.46 | 36.72 | 39.48 |  | 28.76 | 0.63 |
| V4 |  | 36.51±0.62 |  |  | 36.28 | 38.44 |  |  |  |
| V5 | 33.64±1.03 | 36.06±0.60 | 36.07±0.67 | 34.51 | 36.25 | 39.20 |  | 37.94 | 0.53 |
| V6 | 32.83±0.14 | 36.07±0.63 | 36.29±0.68 | 34.64 | 36.15 | 38.79 |  | 36.26 | 0.61 |
| V7 | 33.53±0.00 | 34.62±0.58 | 35.75±0.61 | 33.02 | 35.31 | 38.36 |  | 34.34 | 0.60 |

**Fig. S4.**
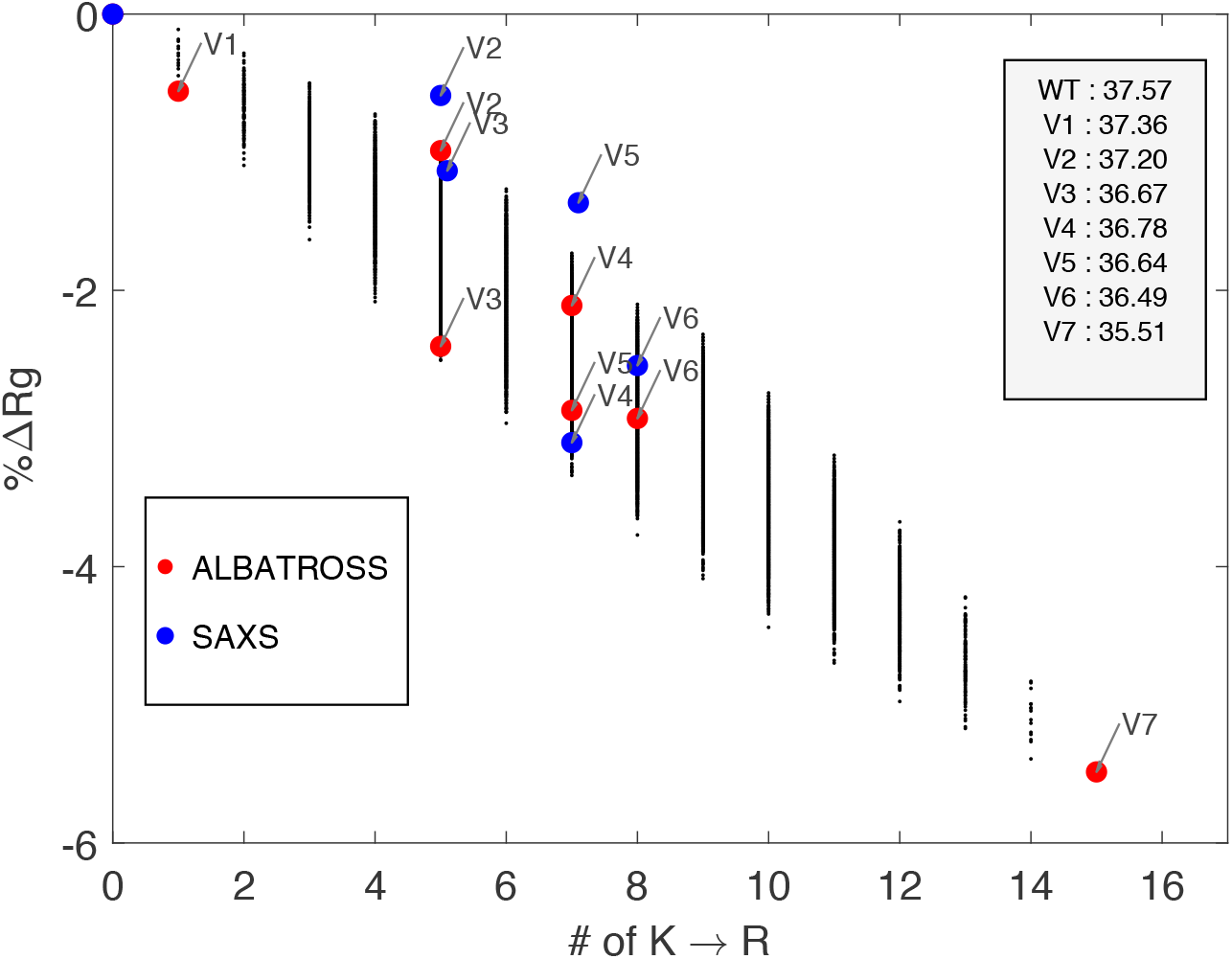
Comparison of *α*Syn *R*_*g*_ changes across mutations at 150 mM NaCl. Black dots show the full ensemble of ALBATROSS simulations (all 7000 models). Red dots highlight specific ALBATROSS mutants of interest. Blue dots indicate SAXS experimental measurements. Arrows and labels mark individual mutants for clarity. The y-axis shows percent change in *R*_*g*_ relative to the WT. The table lists the *R*_*g*_ values predicted by ALBATROSS.

**Fig. S5.**
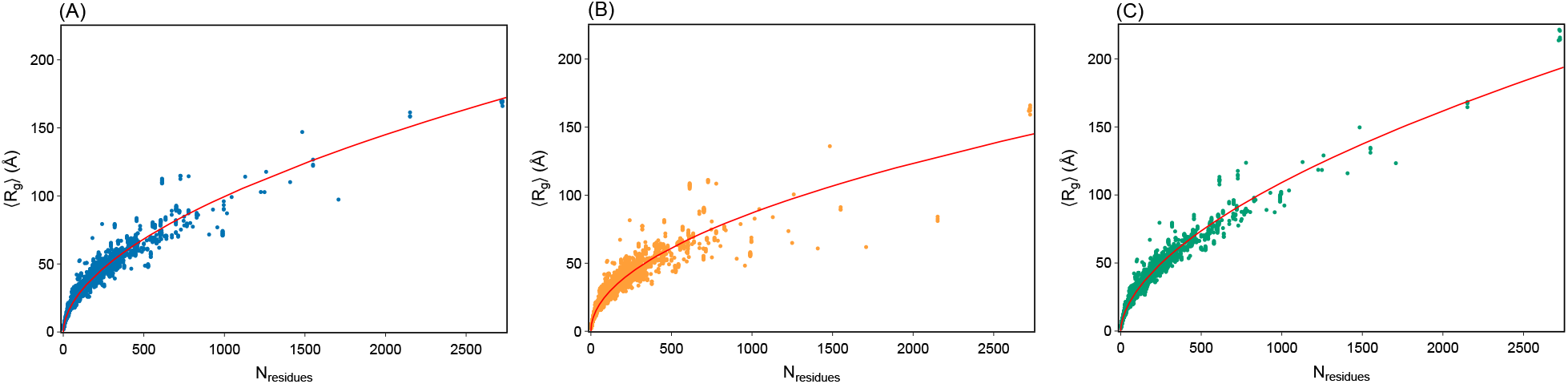
Scaling analysis of DisProt sequences and the structural impact of Arginine-Lysine substitutions. (A) Average radius of gyration (⟨*R*_*g*_⟩) calculated using the CALVADOS force field as a function of sequence length (*N*) for all intrinsically disordered proteins in the DisProt database, where the wild-type (WT) ensemble follows the scaling relationship ⟨*R*_*g*_⟩ = 2.35*N* ^0.54^. (B) The corresponding scaling behavior for the same sequence ensemble following a global lysine-to-arginine (K→R) substitution, resulting in a more compact scaling exponent (⟨*R*_*g*_⟩ = 2.64*N* ^0.51^) due to the enhanced “stickiness” of Arginine residues. (C) Scaling behavior for the ensemble following a global arginine-to-lysine (R→K) substitution, which yields a more expanded state with a higher scaling exponent (⟨*R*_*g*_⟩ = 2.16*N* ^0.57^), reflecting the reduced interactive capacity of the Lysine side chain compared to Arginine. These results demonstrate how the specific identity of cationic residues modulates the global dimensions and polymer scaling properties (*ν*) of disordered protein ensembles while maintaining a constant net charge.

**Fig. S6.**
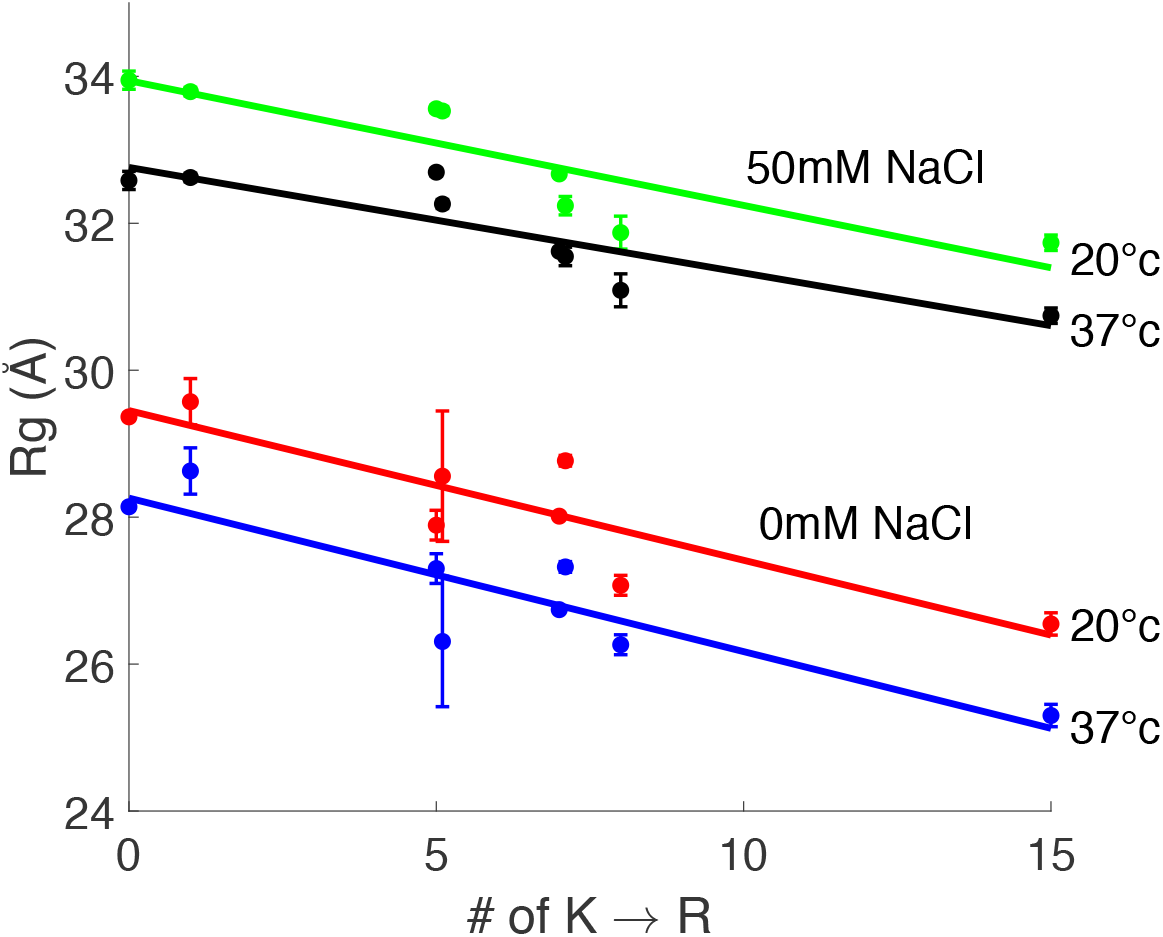
*R*_*g*_ as a function of the number of lysine-to-arginine (K→R) substitutions in *α*-syn. Measurements were performed by SAXS at 20 °C and 37 °C under two salt conditions (0 mM and 50 mM NaCl). Error bars correspond to the uncertainty of the extrapolated Rg. Solid lines indicate linear fits.

**Fig. S7.**
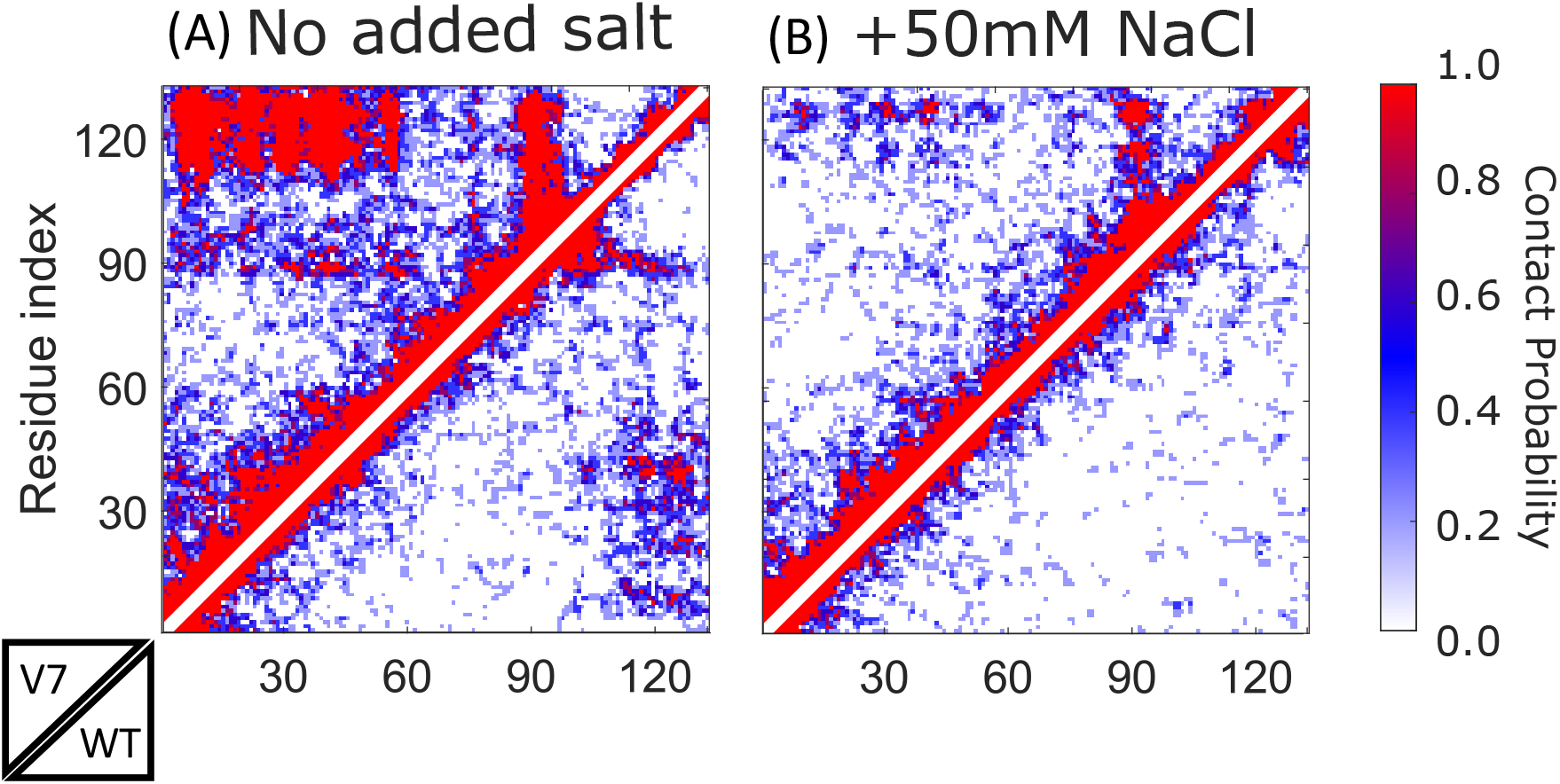
Salt- and K→R-dependent modulation of *α*Syn conformational ensemble. Contact maps from Mpipi-Recharged MD simulations (A,B) show *α*Syn without (A) and with salt (B). In each map, the upper triangle portion represents V7 variant, while the lower portion corresponds to WT. Contacts are normalized to the total number of contacts per ensemble.

**Fig. S8.**
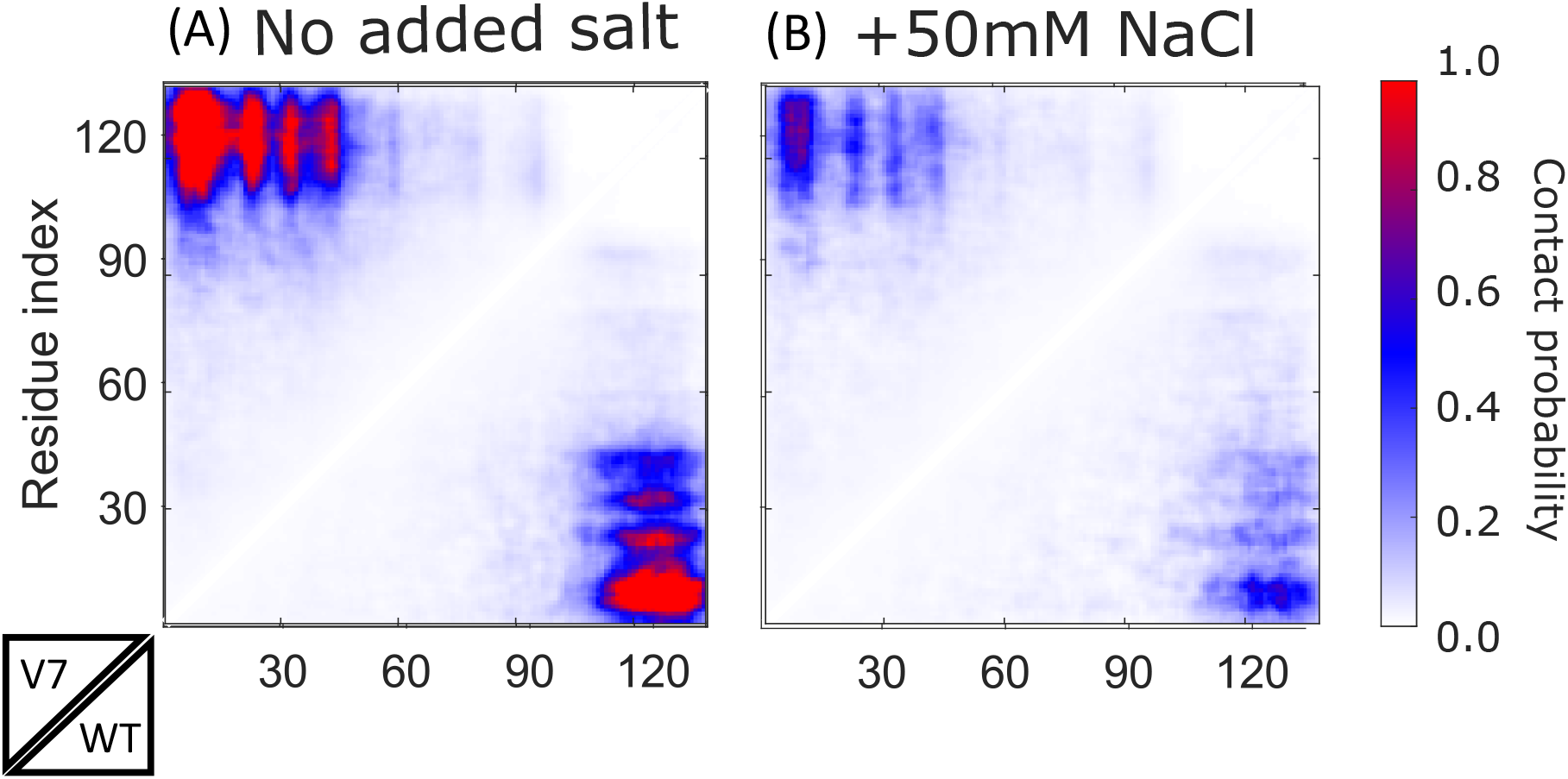
Salt- and K→R-dependent modulation of *α*Syn conformational ensemble. Contact maps from CALVADOS+BME (A,B) show *α*Syn without (A) and with 50 mM NaCl (B). In each map, the upper triangle portion represents V7 variant, while the lower portion corresponds to WT. Contacts are normalized to the excluded volume.

**Fig. S9.**
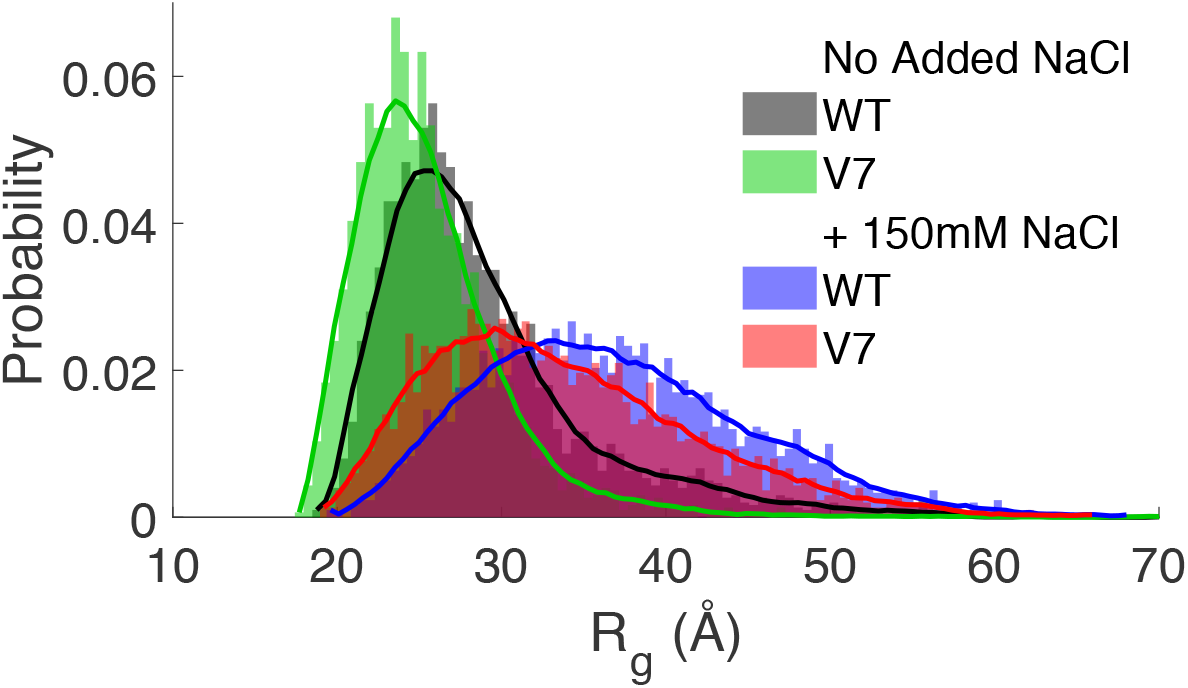
*R*_*g*_ distributions from CALVADOS+BME illustrate ensemble compaction for WT and V7 variant without and with 150mM NaCl

**Fig. S10.**
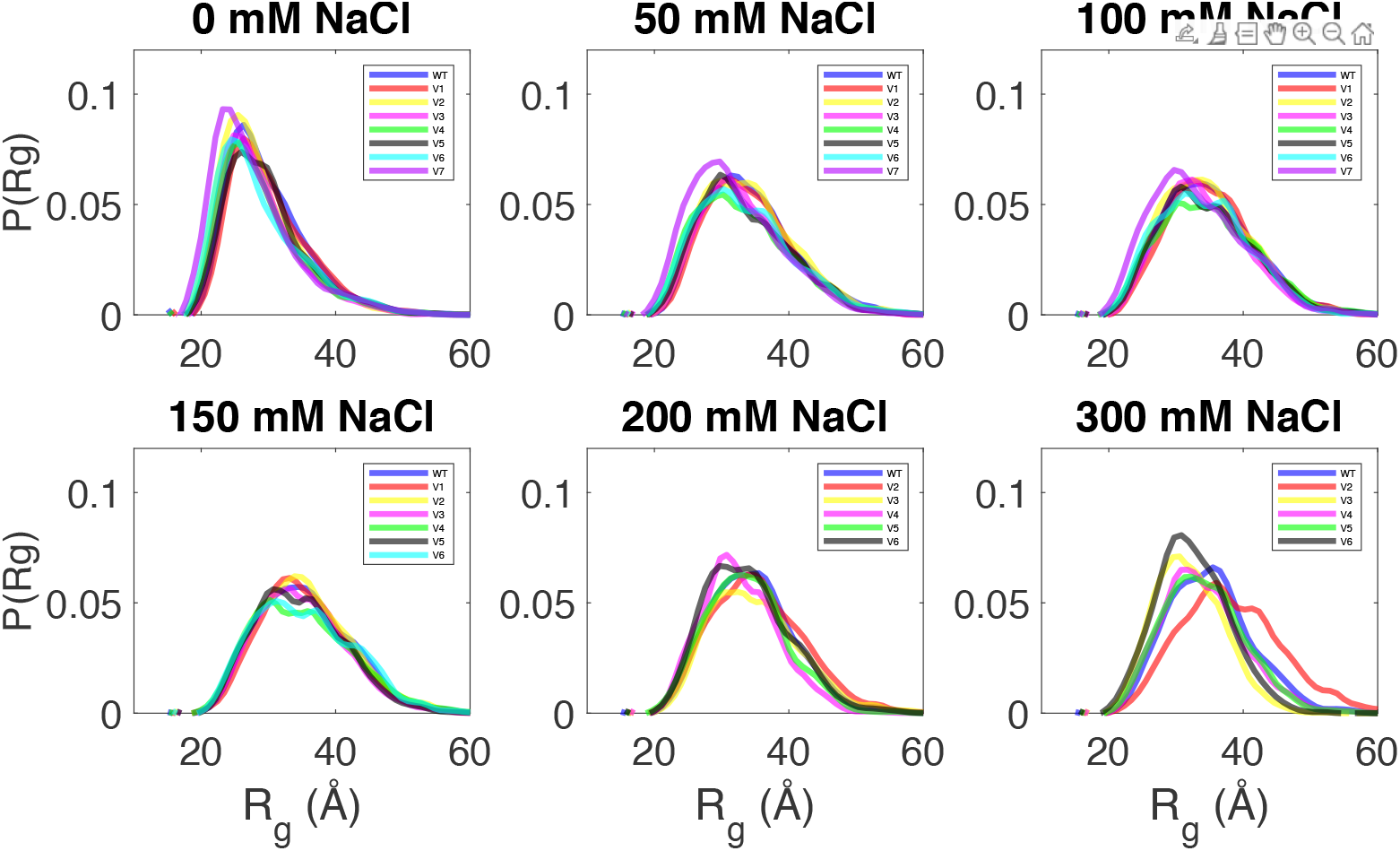
Ensemble Optimization Method (EOM) distributions of *α*-syn variants across a range of ionic strengths. Probability distributions of the *R*_*g*_ for WT and variants V1–V7. Each panel displays the selected conformational ensemble at salt concentrations ranging from 0 to 300 mM NaCl. In all variants, the absence of added salt corresponds to the most compact and narrowest distributions. Increasing salinity (50–300 mM) results in a systematic shift toward larger *R*_*g*_ values and a broadening of the conformational ensemble, indicating increased structural heterogeneity.

**Fig. S11.**
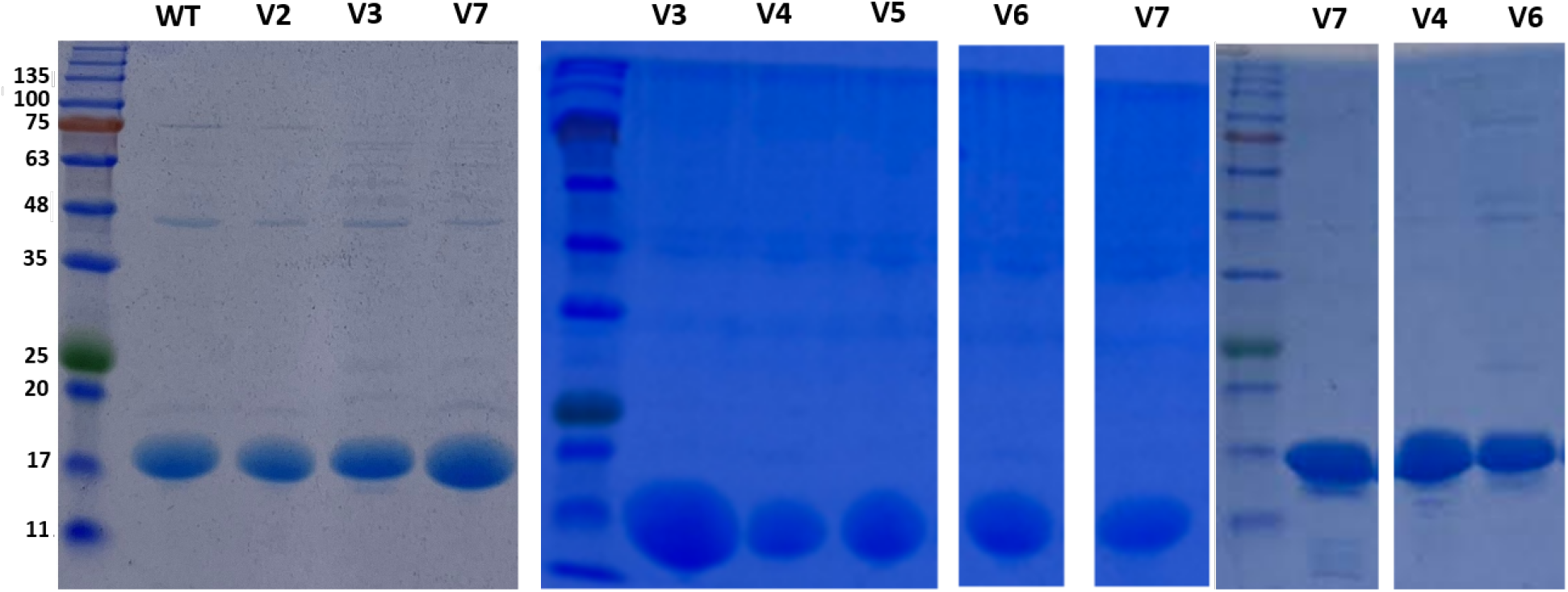
SDS–PAGE analysis of purified WT protein and variants. All samples show a single predominant band at the expected molecular weight with no detectable mobility shift between constructs. Molecular weight markers are indicated in kDa. Although the theoretical molecular weight of *α*-syn is 14.4 kDa, it migrates at ∼17 kDa due to its intrinsically disordered nature. Densitometric analysis confirms >90% purity for all preparations. Additional variants exhibited comparable electrophoretic profiles and purity, indicating consistent expression and purification quality.

**Fig. S12.**
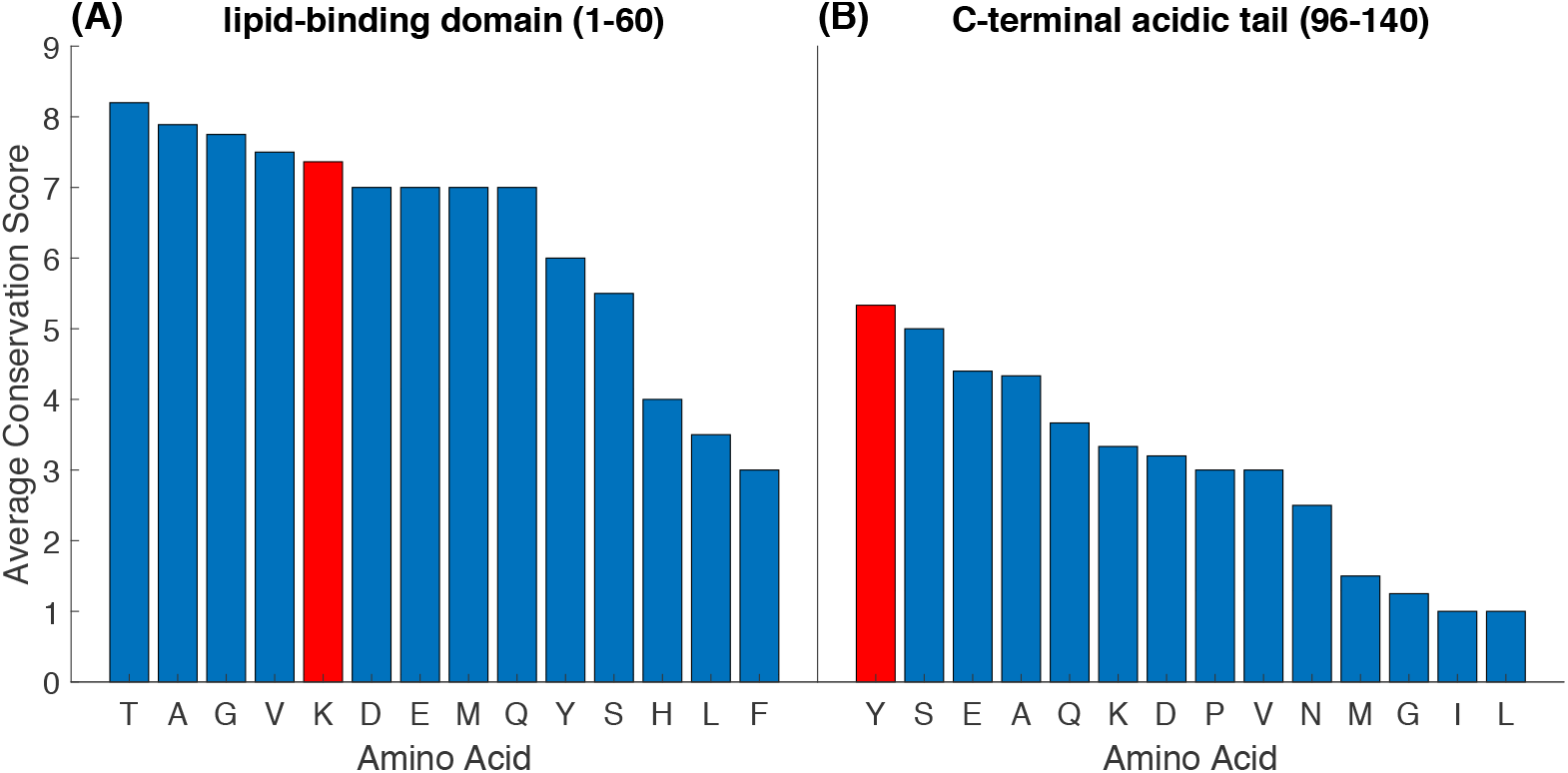
Average amino acid conservation scores calculated using the ConSurf Server (19). (A) Conservation profile of the N-terminal binding domain, with lysine (K) residues highlighted in red. (B) Conservation profile of the C-terminal domain, with tyrosine (Y) residues highlighted in red.

**Fig. S13.**
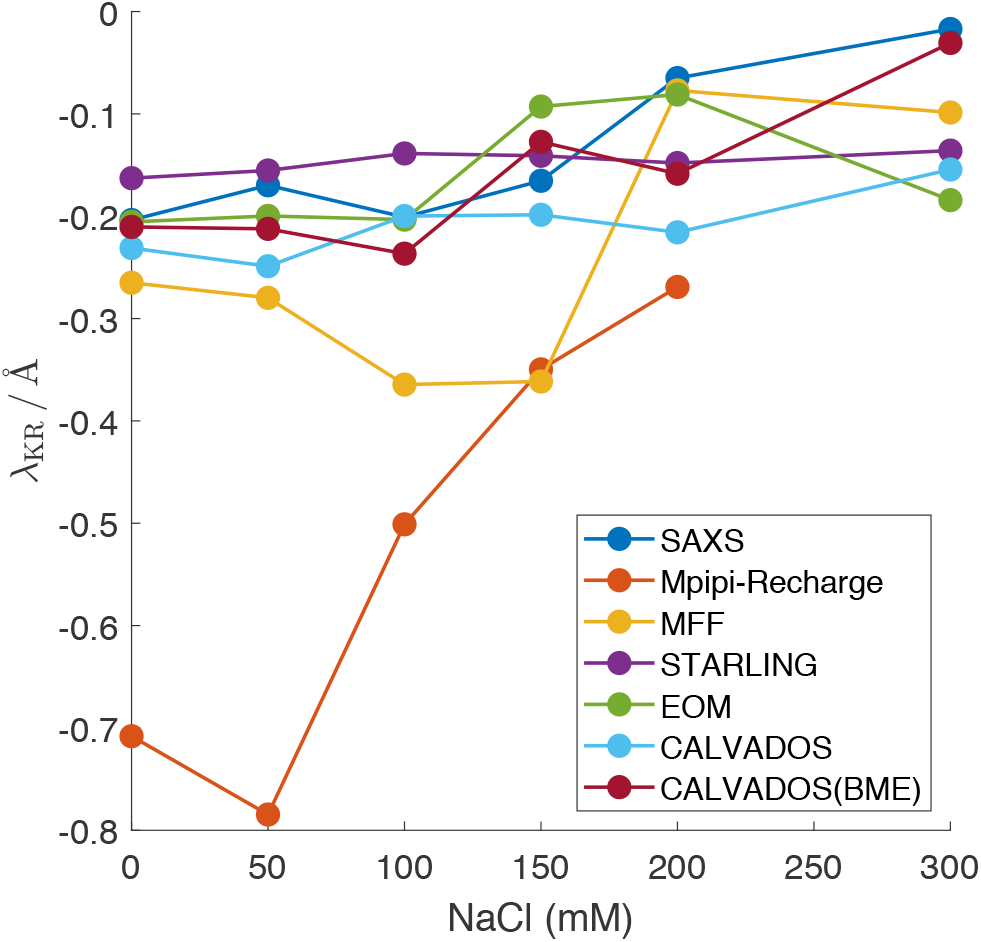
Comparative analysis of the *K* → *R* compaction parameter (*λ*) across experimental and computational methods. The *λ* parameter represents the relative change in chain dimensions resulting from *K* → *R* substitutions. Experimental *λ* values derived from SAXS measurements as a function of ionic strength, showing a persistent compaction effect up to 150 mM NaCl followed by a collapse toward baseline at 300 mM NaCl. Comparison of *λ* values calculated from various computational frameworks and force fields, including Mpipi-Recharge, MFF, STARLING, EOM, and CALVADOS.

